# Mass mortality of southern elephant seals during multi-species outbreak of HPAI H5N1 on sub-Antarctic Heard Island

**DOI:** 10.64898/2026.06.16.732752

**Authors:** Julie C. McInnes, Tristan Burgess, Georgia Mergard, Melanie R. Wells, Clive R. McMahon, Matthew J. Neave, Andrea Polanowski, Aleks Terauds, Jérémy Tornos, Mathilde Lejeune, François-Xavier Briand, Guy Baele, Thierry Boulinier, Helen Achurch, Rachael Alderman, Anna Lashko, Barbara Wienecke, Louise P. Wynen, Benjamin Viola, Patti Virtue, Jarrod C. Hodgson

**Affiliations:** Australian Antarctic Division, Department of Climate Change, Energy, the Environment and Water, Kingston, Tasmania 7050, Australia; Institute for Marine and Antarctic Studies, University of Tasmania, Battery Point, Hobart 7004, Australia; School of Infectious Diseases and Global Health, Tufts University’s Cummings School of Veterinary Medicine North Grafton, MA, 01536, USA; Center for Wildlife Studies, Camden, ME 04843 USA; IMOS Animal Tagging, Sydney Institute of Marine Science, 19 Chowder Bay Road, Mosman, 2088, New South Wales, Australia; Australian Centre for Disease Preparedness, Commonwealth Scientific and Industrial Research Organisation (CSIRO), Geelong, VIC 3220, Australia; Centre d’Ecologie Fonctionnelle et Evolutive (CEFE), CNRS, Université Montpellier, EPHE, IRD, Montpellier, France; ANSES, Ploufragan-Plouzané-Niort Laboratory, National Reference Laboratory for Avian Influenza, Unité de virologie, Immunologie, Parasitologie, Aviaires et Cunicoles, Ploufragan, France; Department of Microbiology, Immunology and Transplantation, Rega Institute, KU Leuven, Leuven, Belgium

**Keywords:** *Mirounga leonina*, Southern Ocean, high pathogenicity avian influenza, HPAI phylogeography, transmission patterns

## Abstract

High pathogenicity avian influenza (HPAI) has spread across the sub-Antarctic, causing significant wildlife impacts. We report its first detection in an Australian external territory, Heard Island and McDonald Islands, which supports over one million breeding seabirds and seals. Drone and ground surveys (October 2025, January 2026), combined with viral genome analysis, confirmed infection with Influenza A H5N1 clade 2.3.4.4b at Heard Island. Drone surveys revealed mass mortality in southern elephant seals, with 8,573 pups (62%) recorded dead across Heard Island by the final surveys. Mortality increased at an average rate of 5.6% per day in a subset of harems, and the highest observed mortality in a harem was 97%. Based on the average (76%) mortality in the final surveys, total estimated pup mortality at Heard Island was 13,359 (from a total population of 17,364 pups), though this may be an underestimate as mortality was ongoing at this time. HPAI was detected in six of nine species tested and, we suspect, led to elevated mortality in king and gentoo penguins. Phylogenetic analysis indicates the virus was introduced from Crozet Islands, with an estimated arrival around August 2025. These data show the continued easterly spread of HPAI around the sub-Antarctic, with severe but heterogeneous impacts across taxa. Our results demonstrate the value of drones for large-scale monitoring, underscoring the need for continued and enhanced HPAI surveillance across the Southern Ocean.

## INTRODUCTION

High pathogenicity avian influenza (HPAI) strain H5N1 clade 2.3.4.4b has caused a global panzootic since 2021. Wild birds and mammals on all continents other than Australia have been affected^1,2^. In 2022, the virus spread around the South American coastline causing widespread mortality in seabirds and seals^3,4^. During this period new mutations were evident, some of which are thought to facilitate mammal-to-mammal transmission^5^, accompanied by increasing mass mortalities in pinnipeds^2^. In spring 2023, the virus spread from the southern cone of South America to the Falkland Islands/Islas Malvinas, South Georgia Island/Islas Georgias del Sur^6–8^ and then the Antarctic Peninsula^9–11^. The virus continued to spread in 2024, moving from South Georgia towards the Indian Ocean basin, and was detected on Gough Island, Prince Edwards Islands (on Marion Island), Crozet Islands (on Possession Island) and Kerguelen Islands in September, October and November, respectively^12–14^. There is no evidence of HPAI H5N1 on mainland Australia (including Tasmania), Macquarie Island, New Zealand, the New Zealand sub-Antarctic islands^15^ or the Antarctic continent outside of the Antarctic Peninsula region ^16,17^.

The variant of the H5N1 virus that had reached southernmost South America in 2023 and spread to the adjacent Antarctic and sub-Antarctic islands is distinct from variants circulating in Europe and North America. All viral segments are avian in origin, containing a mixture of Eurasian and north American genetic lineages. The progenitor viruses have reassorted extensively in North America in wild birds^18^. The viruses found in South America and the Antarctic belong to genotype B3.2 and derive from a single introduction from North America^18,19^. This clade acquired several PB2 mutations likely to facilitate mammalian transmission^13^. Recent analysis suggests the development of two distinct genetic sub-lineages within clade 2.3.4.4b in the south Atlantic: “Clade I”, which has spread eastward from South Georgia to sub-Antarctic islands and derived primarily from viruses that circulated in birds in South America; and “Clade II”, which circulated primarily in marine mammals in South America and has continued to circulate in the Falklands/Islas Malvinas, South Georgia/Islas Georgias del Sur and the Antarctic Peninsula^13^. Neither clade, however, appears to be specific to either mammals or birds^13^, and phylogenetic analysis suggests that the virus was introduced independently to the Kerguelen and Crozet islands from South Georgia during 2024^12^.

The impact of HPAI on Antarctic and sub-Antarctic species is concerning due to high-density aggregations of marine mammals and birds, low species diversity, a high degree of endemism and the predominance of long-lived species with low reproductive output (i.e. *k-*selected species), many of which are globally threatened and/or have declining populations. High mortality rates in adult animals can have a considerable impact on long-term population trajectories, while mortality of offspring can have lasting impacts on populations if the virus persists in a region for multiple years^20^. Unusual mortalities caused by HPAI have been observed as animals return to breed, peaking at the height of the breeding season^6^. This likely reflects the return of infected individuals from migration to breeding colonies, where animals congregate in high densities. Observer bias may also contribute to this pattern, as breeding colonies are generally accessible to researchers to conduct observations and testing, whereas mortality at sea cannot be easily detected. Incursion estimates based on genomic data from the Antarctic Peninsula, South Atlantic and southern Indian oceans support the winter/early spring colony returns as a key time window for introductions^13^.

The impact of HPAI has varied across sub-Antarctic mammal and bird species. The most severe impact has been on southern elephant seals (*Mirounga leonina*) with mass mortality events observed at Península Valdés, Argentina and South Georgia/Islas Georgias del Sur^21,22^. At Península Valdés, 96% pup mortality was observed by the end of the 2022 breeding season, with >17,000 pups predicted to have died^20^. The population consequences were still apparent a year later, when only one third of the individuals normally expected to breed returned^21^. At South Georgia, there was a 47% decrease in the number of breeding females between 2022 and 2024 at the three largest breeding colonies^21^. In both cases, ongoing monitoring will be required to understand whether this is reflective of high adult mortality or skipped breeding. High mortality has been reported in brown and south polar skuas (*Stercorarius antarcticus and S. maccormicki*)^6,10,11^, snowy (*Diomedea exulans*) and black-browed albatross (*Thalassarche melanophris*)^6,8,23^, king (*Aptenodytes patagonicus*) and gentoo (*Pygoscelis papua)* penguins^12,24^, and variable mortality in Antarctic fur seals, but this is difficult to quantify and may be much higher than reported^6,13^. Some species, such as southern giant petrel (*Macronectes giganteus)* have been surveyed in several areas with no unusual mortality reported^6, 10^. Accurately assessing mortality rates remains challenging due to spatial and temporal constraints on site access. Drone-facilitated surveys enable broad-scale monitoring, overcoming some of these constraints. When combined with ground sampling, drone surveys can substantially improve HPAI impact assessments^25^.

Heard Island and McDonald Islands (HIMI), located 4,000 km south-west of Australia, are home to over one million breeding seabirds and seals that have not been surveyed in over 20 years. Nineteen species of seabird and three seal species breed on the islands, including two endemics: the Heard Island shag (*Leucocarbo nivalis)* and a subspecies of black-faced sheathbill (*Chionis minor nasicornis*). The islands are home to globally important populations of king penguins, southern elephant seals, southern giant petrels, macaroni penguins (*Eudyptes chrysolophus)* and the largest populations of black-browed albatross and gentoo penguins in the Australian territory. HIMI is located <450 km south-east of the Kerguelen Islands and <1,700 km from the Crozet Islands and animals are known to move between these sites^26–28^.

The Australian Antarctic Program conducted two voyages to HIMI in October 2025 and January 2026. We conducted surveillance for HPAI and tested the assumption that HPAI had spread from neighbouring islands. Using drone and ground surveys, together with multi-species sample collection, we investigated the spatial and temporal distribution of mortality across mammal and bird species and analysed the spread in elephant seals. Here, we report on the unusual mortality of seabirds and seals at HIMI documented during those surveys. Samples from carcasses facilitated the sequencing of the HPAI virus, providing novel insights into the phylogeographic nature of the circumpolar spread.

## RESULTS

Over the two voyages, 120 drone flights were completed totalling more than 54 hours of airtime and approximately 1,600 km of distance flown. Nearly all flights extended beyond visual line of sight (BVLOS; 98%) and the majority were conducted from the ship (RSV *Nuyina*; 64%). Some flights were mapping missions used to produce approximately 16 km^2^ of high-resolution orthomosaics. Ground searches covered approximately 8.8 km^2^ (Fig. 1).

**Fig. 1:**
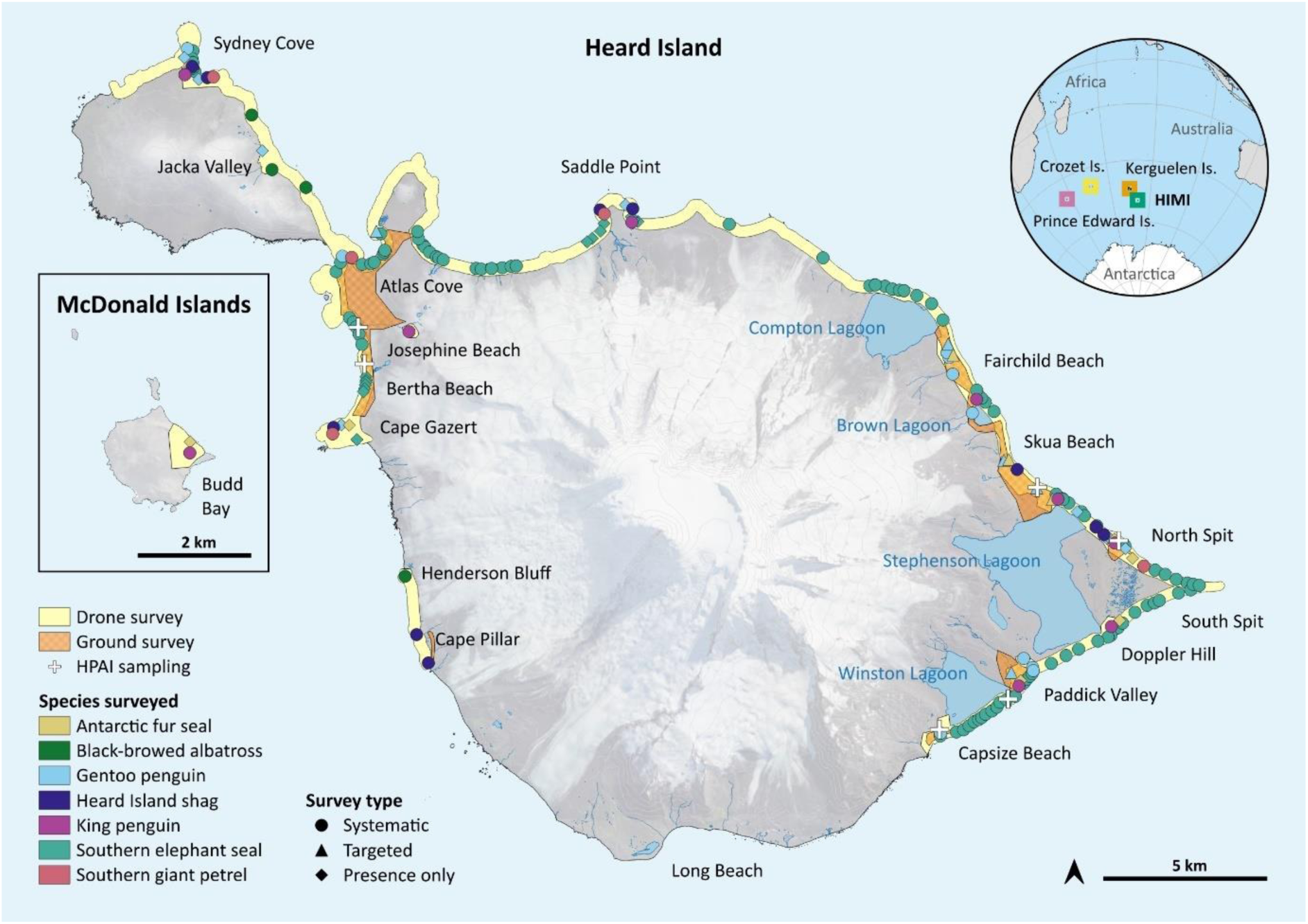
Survey and sampling effort for the presence of HPAI in a subset of seabird and seal species (n = 7) across Heard Island and McDonald Islands (HIMI) in October 2025 and January 2026. Drone surveys covered most of the ice-free areas of the coast of Heard Island and a portion of McDonald Island, with ground surveys conducted in a subset of areas on Heard Island. All surveys were either systematic or targeted. Some colonies are shown as presence only when they were identified but not surveyed for HPAI.

### Systematic counts

The entire northern coastline of Heard Island was surveyed for elephant seals from 12–22 October 2025 and along sections of the southern coastline from 16–22 October (Fig. 1). Across all areas, 109 harems were identified: 87% (n = 95) were surveyed using drones enabling abundance estimates of pups and adults, and the remainder (n = 14) were surveyed using helicopters, enabling counts of adults only. Unusual mortality was observed via drone and ground surveys (Fig. 2). The total number of pups counted from drone imagery was 13,883, with 8,573 (62%) of those not within the normal close proximity of an adult female (hereafter unpaired; Fig. 3) and thus presumed dead or moribund. Pup mortality within harems ranged from <1% at the start of the survey period in the north-west of the island to 97% at the end of the survey period at Winston Lagoon in the south-east (Fig. 4). There was a strong correlation between mortality and proximity to Winston Lagoon and observed pup mortality reduced with greater distance from the region (Fig. 4; Fig. S2; Table S3). Repeat surveys of 16 harems at the eastern end of the island, along the northern (October 13, 16, 22) and southern (October 16, 21, 22) sides of the Elephant Spit, revealed an average daily increase in pup mortality of 5.6% (range 0.1–13.3% per day). This rate was highest furthest from Winston Lagoon (Fig. 4c), suggesting pup mortality may have plateaued around Winston Lagoon by this time, while other areas were likely still in the exponential increase phase of the epidemic curve. Pup mortality on the final survey dates (21–22 October) averaged 76% (range 53–97%). As surveys from 20–22 October were restricted to the eastern end of the island where the highest mortality was seen, separating spatial and temporal drivers of mortality is difficult using October data alone. However, in January 2026 widespread elephant seal mortality was evident across all known breeding areas, confirming the disease had spread island-wide, affecting the entire breeding population. Although the carcasses were decomposing by this time, the dead pups were small and still covered in their natal coat, indicating that they were not yet weaned (Fig. 2). Given the highly synchronous breeding behaviour of elephant seals and rapid growth rates of pups, this indicates HPAI spread around the island within two weeks of the October surveys, suggesting mortality was likely high in all areas by 20–22 October 2025.

**Fig. 2:**
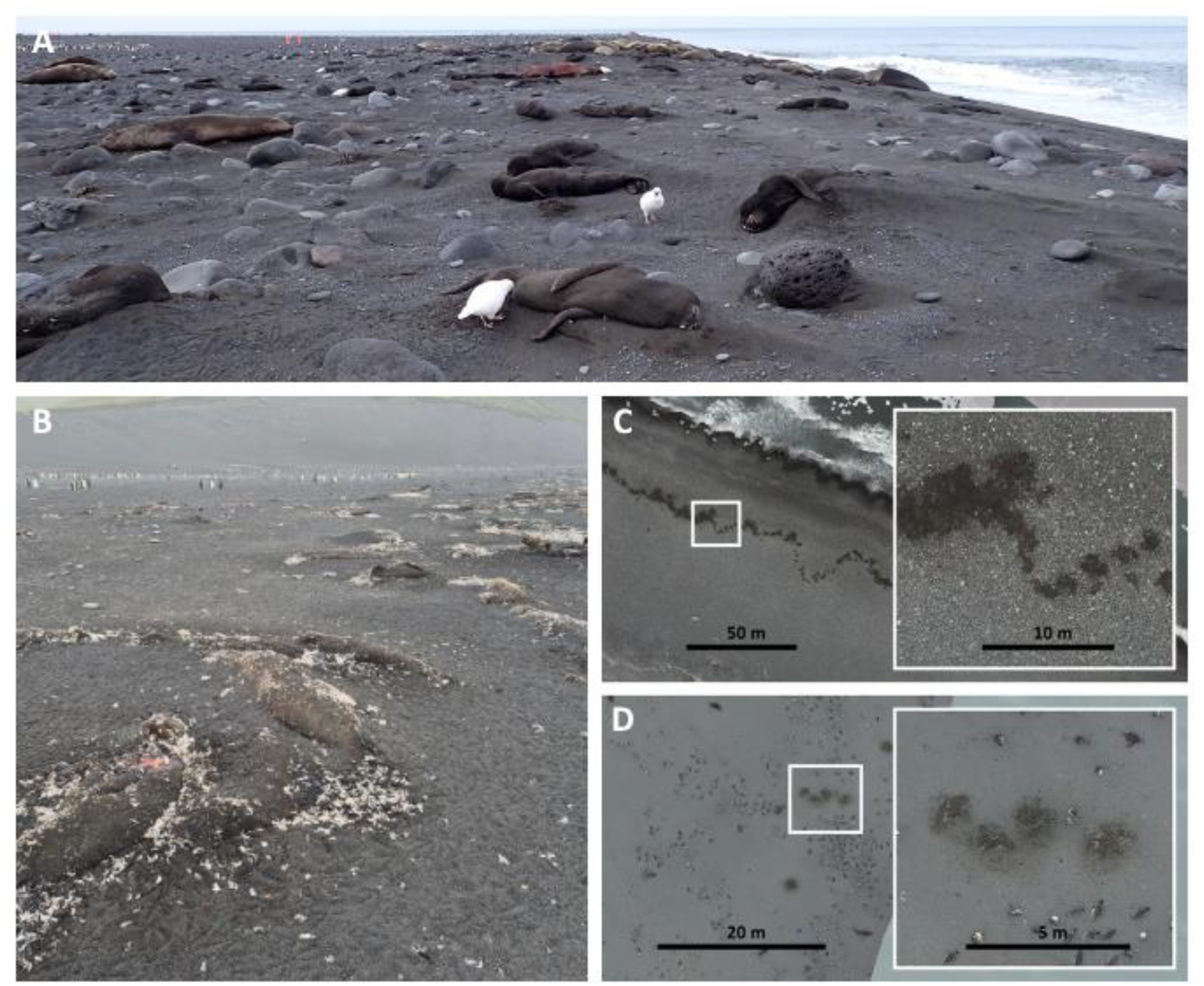
Southern elephant seal pup mortality and carcass decomposition in October 2025 and January 2026 at Heard Island and McDonald Islands. **(A)** Unusual mortality was seen at Winston Lagoon Beach during October 2025. Note scavenging on pup carcasses by black-faced sheathbills *(Chionis minor nasicornis*). In January 2026, there was evidence of widespread pup mortality, with variability in carcass decomposition, around all breeding areas covered by **(B)** ground surveys at Josephine Beach, Heard Island, and drone surveys at **(C)** northern Elephant Spit, Heard Island and **(D)** Budd Bay, McDonald Island.

**Fig. 3:**
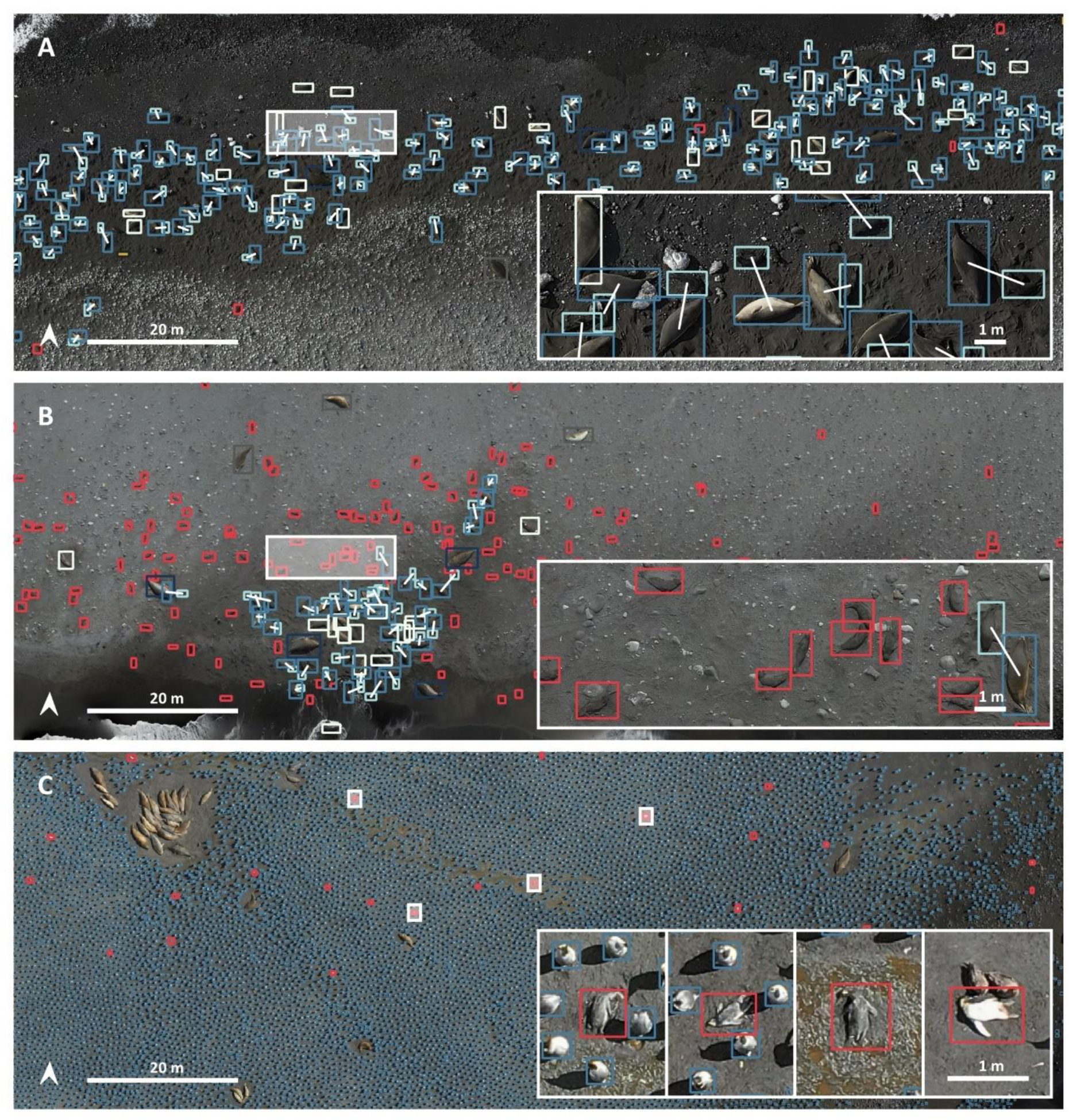
Drone imagery was processed into high-resolution orthomosaics to estimate the impact of HPAI. **(A)** Southern elephant seals were categorised using spatial proximity rules (red rectangles = unpaired pup, blue rectangles = cohorts of live animals (see Table S1), white lines = paired pups with adult females; see Supplementary Methods for more detail) at Compton Lagoon, Heard Island on 14 October 2025. **(B)** Elephant seals at Winston Lagoon, Heard Island on 21 October 2025. **(C)** King penguin breeding colony at Doppler Hill, Heard Island on 4 January 2026. Fresh carcasses (red rectangles) were manually annotated, while the total number of adult penguins (blue rectangles) were predicted using a bespoke YOLOv11 model.

**Fig. 4:**
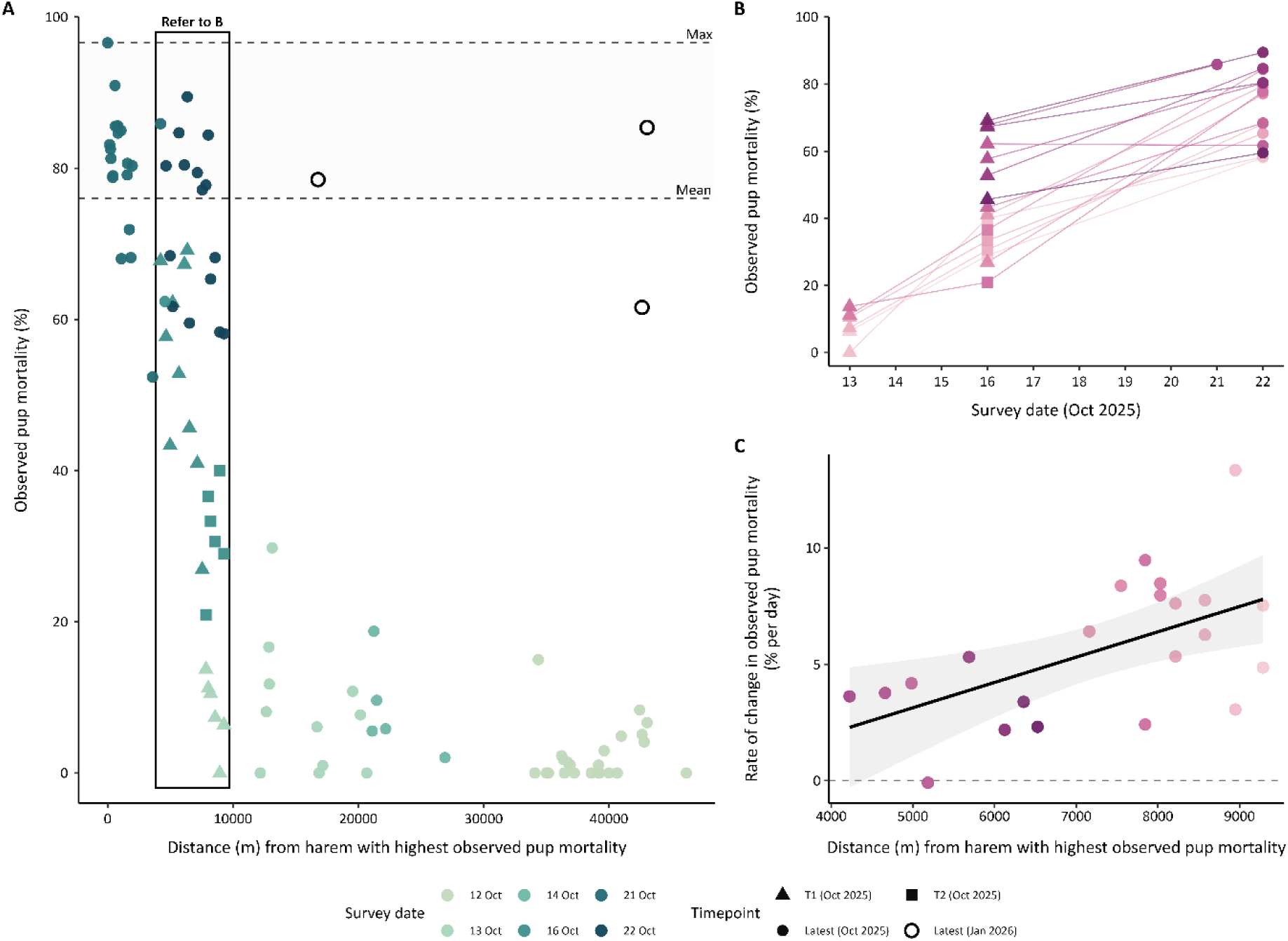
Observed southern elephant seal pup mortality at Heard Island in October 2025 and January 2026. **(A)** Observed mortality across harems (with ≥10 pups) surveyed by drone (n = 74) relative to their distance from Winston Lagoon, where the highest pup mortality was detected, noting surveys furthest from Winston Lagoon occurred earlier in October 2025 than those in close proximity. The maximum and mean observed pup mortality for the latest October 2025 surveys are shown. In January 2026, three locations (hollow circles) were ground-surveyed to estimate pup mortality (Brown Lagoon north = 2 harems; Josephine Beach north = 3 harems; Josephine Beach south = 2 harems). **(B)** Increase in pup mortality over time derived from a subset of harems that were surveyed multiple times during October (n = 16; box in **A**, pink/purple symbols in **B** and **C**. **(C)** Daily mortality rate estimated from harem mortality in B relative to distance from Winston Lagoon.

Ground counts were completed along a 1.4 km section of the coast at Winston Lagoon. On 21 October 2025, 1,189 pups were counted, of which 88% were dead and another 18 were showing clinical signs of HPAI, including laboured breathing, tremors and twitching, marked incoordination, unresponsiveness and lethargy. These data support our drone-facilitated mortality estimates and indicated some females were still attending pups or remained with dead pups, suggesting mortality estimates derived from drone surveys alone were likely an underestimate (Fig. S1; Table S2). On this section of the coast, nine dead female elephant seals were seen representing 3% of the 271 breeding females counted.

During October, we estimated pup production at 17,364, which likely underestimates total births as parturition was still ongoing in some areas at the time of surveys. Using this conservative estimate and the mortality rates recorded at the final surveys, the total number of dead pups by this time was estimated to be 13,359. However, if the highest observed mortality (97%) occurred at all harems, pup mortality may have been closer to 16,791.

Although neonate mortality during the October 2025 survey at Josephine Beach and Brown Lagoon was low, 5.0% and 5.8% respectively, carcass counts in January were 61.6% and 85.4% at north and south Josephine beaches, respectively, and 78.5% mortality at Brown Lagoon (Fig. 4, Table S8). These estimates are approximate as beaches are dynamic, and it was highly likely that some carcasses had washed away. The counts from January 2026 highlight the overall substantial pup mortality across Heard Island coupled with evidence of mortality at McDonald Island (Fig. 2).

Elevated levels of mortality were detected in king penguins using data collected in January 2026 (Fig. 3). Of the 235,000 adult king penguins estimated at and around breeding colonies through systematic counts, 298 freshly dead and suspected dead adults were counted (Table S7). While the proportion of dead individuals was low (<1% at all sites), it is unusual to find dead adult king penguins, even in large colonies (compared with 3 dead/ ∼110,000 adults at Macquarie Island, Supplementary Methods). Most of the carcasses (n = 253; 85%) were detected in the south-east of Heard Island at Doppler Hill, the largest colony. Carcasses were spread through the colony and its periphery, rather than concentrated together (Fig. 3, Fig. S6). Other areas with elevated mortality were Paddick Valley

(n = 12) and McDonald Island (n = 18), compared to <5 dead adults at other colonies. An additional 16 dead adult birds were observed outside breeding colonies during targeted and opportunistic ground surveys, predominantly along Josephine and Bertha beaches; these were old, desiccated carcasses, suggesting prior mortality events. One of the 19 scat samples collected from apparently healthy adults tested positive for HPAI (Table 1). No unusual adult mortalities were detected during October 2025. Five king penguin carcasses were counted in drone imagery, all at Doppler Hill (no surveys were conducted at McDonald Island at this time). One additional dead adult was found during ground surveys along the northern Elephant Spit, Heard Island during October 2025, and one adult and four chicks were sampled. All tested negative for HPAI (Table 1).

**Table 1:**
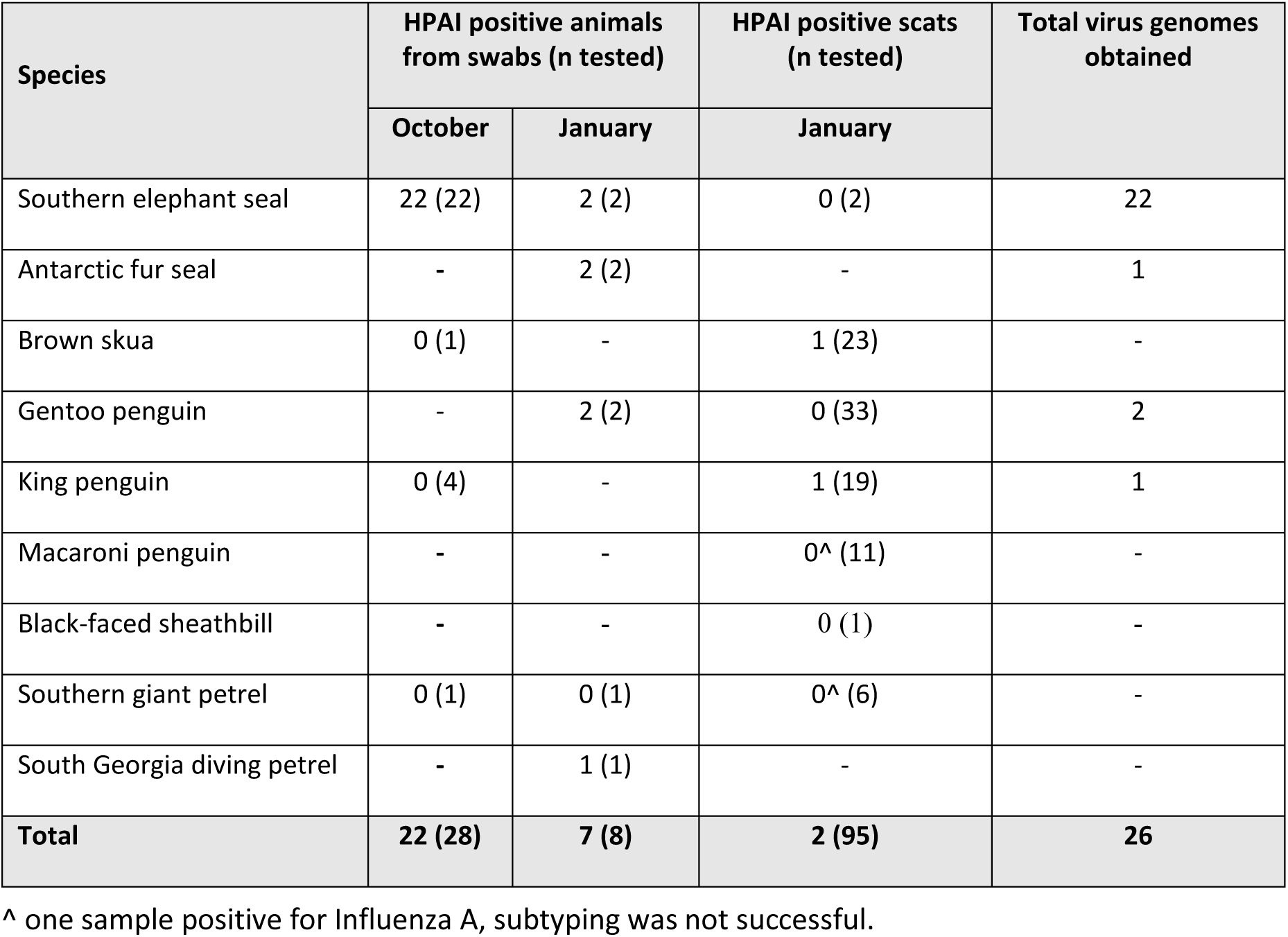
Summary of samples collected at Heard Island in October 2025 and January 2026 that were tested for Influenza A, HPAI and sequenced. All swab samples were collected from animals found dead: where possible, from the oropharynx, cloaca and brain for birds and oral cavity, nose and brain for seals. Fresh scat samples were collected from live, apparently healthy animals.

| Species | HPAI positive animals from swabs (n tested) |  | HPAI positive scats (n tested) | Total virus genomes obtained |
| --- | --- | --- | --- | --- |
|  | October | January | January |  |
| Southern elephant seal | 22 (22) | 2 (2) | 0 (2) | 22 |
| Antarctic fur seal | - | 2 (2) | - | 1 |
| Brown skua | 0 (1) | - | 1 (23) | - |
| Gentoo penguin | - | 2 (2) | 0 (33) | 2 |
| King penguin | 0 (4) | - | 1 (19) | 1 |
| Macaroni penguin | - | - | 0<sup>^</sup> (11) | - |
| Black-faced sheathbill | - | - | 0 (1) | - |
| Southern giant petrel | 0 (1) | 0 (1) | 0<sup>^</sup> (6) | - |
| South Georgia diving petrel | - | 1 (1) | - | - |
| **Total** | **22 (28)** | **7 (8)** | **2 (95)** | **26** |

<sup>^</sup> one sample positive for Influenza A, subtyping was not successful.

On Heard Island, 24 dead adult gentoo penguins were seen in January 2026: 21 detected through systematic counts of camera imagery (5,125 live adults) and three additional animals were observed during ground surveys. While 13 dead birds at Brown Lagoon were seen in both ground and drone imagery, they have only been allocated to the drone counts to avoid duplication. These carcasses were showing some decay suggesting mortalities prior to January. Swab samples were obtained from two fresh carcasses and tested positive for H5N1 (Table 1).

No unusual mortalities were detected in any other species on Heard Island (Table S5, S6 for counts of carcasses). All known breeding areas of black-browed albatross, shags and ∼20% of southern giant petrel colonies were systematically searched (Fig. 1), and ground surveys were conducted in the highest-density breeding area of Antarctic fur seals (representing >50% of the island population). One dead black-browed albatross and one bird with a droopy wing were seen at the Henderson Bluff and Jacka Valley colonies, respectively (Table S6; Fig. S7). Four southern giant petrels, three brown skuas, six adult fur seals (plus six pups), three macaroni penguins and one South Georgia diving petrel *(Pelecanoides georgicus)* were found dead (Table S5) but only a subset could be sampled (Table 1).

### Influenza A molecular testing

Swab samples were collected from 36 dead animals, 28 in October 2025 and eight in January 2026 (Table 1). Of these, 29 individuals (80%) tested positive for both Influenza A and H5N1. This included all elephant seals (n = 24), Antarctic fur seals (n = 2), gentoo penguins (n = 2) and the single South Georgia diving petrel. To our knowledge, this is the first report of HPAI mortality in a diving petrel with a positive brain swab. Of these, whole virus genomes could be obtained for 24 samples, and all positives were type H5N1 clade 2.3.4.4b. Brain swabs consistently returned lower qPCR cycle threshold scores and were therefore prioritised for high-throughput sequencing (Supplementary Data). Scat samples were collected from 95 individuals across seven species. Of these, one king penguin and one skua tested positive for HPAI, while one southern giant petrel and one macaroni penguin tested positive for Influenza A, but subtyping was unsuccessful on these samples (Table 1).

### Phylogenetic placement and molecular dating of Heard Island H5N1 viruses

The Heard Island H5N1 2.3.4.4b viruses fell within the Antarctic/sub-Antarctic sub-lineage designated Clade I by Clessin et al.^13^ (see their Fig. 2). This places them adjacent to viruses that spread eastward around the Southern Ocean to Crozet, Kerguelen, Prince Edward and Gough islands^13^. All Heard Island sequences were classified as genotype B3.2, consistent with other Antarctic and sub-Antarctic H5N1 2.3.4.4b viruses. Within Clade I, the Heard Island viruses formed a closely related monophyletic cluster in the concatenated whole-genome phylogeny (Fig. 5). This cluster grouped closest to viruses from the Crozet Islands (100% posterior support), which was estimated as the most likely introduction source (100% ancestral location support) through Bayesian phylogeographic inference.

**Fig. 5:**
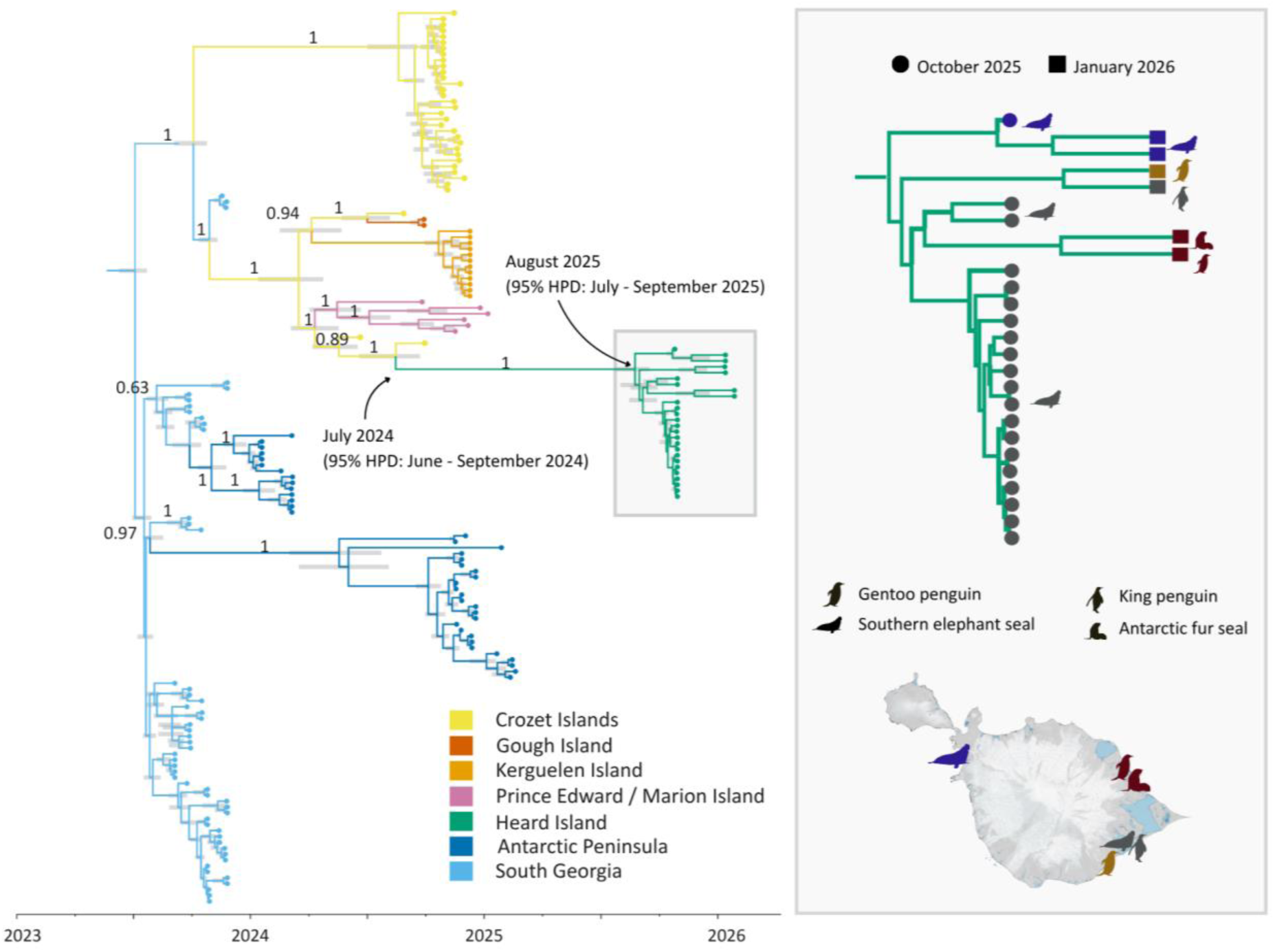
Spatiotemporal reconstruction of HPAI H5N1 2.3.4.4b Clade I dispersal in the sub-Antarctic and Antarctica using an asymmetric forward-in-time discrete phylogeographic model. The source of an estimated single introduction onto Heard Island was estimated to be the Crozet Islands, with 100% ancestral location support. Posterior support values for phylogenetic clustering are provided on the branches of the majority-rule HIPSTR consensus phylogeny, and the node bars indicate the 95% highest posterior density (HPD) intervals on the estimated node ages. The Heard Island infections constitute a monophyletic cluster with a time to most recent common ancestor (tMRCA) in August 2025 (95% HPD: July-September 2025), and a tMRCA with the most closely related Crozet Islands infection in July 2024 (95% HPD: June-September 2024). Within the Heard Island cluster, infections showed closer similarity with sampling location rather than by species or sampling time (see inset).

Molecular-clock analysis estimated the time to the most recent common ancestor (tMRCA) of the sampled Heard Island viruses as August 2025 (95% HPD: July–September 2025), only months before sample collection. However, because the sampled genomes may not capture the full genetic diversity present on Heard Island, the introduction date and the tMRCA of the viruses on Heard Island may have been earlier. The branch separating the Heard Island clade from closest sampled relatives represented approximately one year of evolutionary time (Fig. 5), indicating a substantial period of unsampled evolutionary history before this detection on Heard Island. Root-to-tip regression analysis showed that the Heard Island viruses had lower genetic divergence than expected from their sampling dates, relative to the broader H5N1 2.3.4.4b dataset (Fig. S9).

### Molecular marker characteristics of HIMI H5N1 viruses

The FluMut tool identified 62 mutations across the HIMI dataset. These mutations matched those found in other sub-Antarctic island viruses with only three exceptions: two mutations were found in two elephant seal individuals: a polymerase acidic (PA) R266L mutation in sample HIMI-543, and a non-structural protein 1 (NS1) E55K substitution in sample HIMI-548, which has been associated with interspecies host adaptations^29^. Additionally, a PB2 R251K mutation, that could support interspecies host adaptations^5,12,13,30^ was fixed across the entire HIMI dataset when evaluated against the reference sequences and sub-Antarctic variants (Supplementary Data). Several of the identified mutations are recognized as key markers for specific pathogenic phenotypes^5,12,13,30^. The polymerase basic 2 (PB2) E627K mutation, regarded as the most important marker for mammalian adaptation^31,32^, was notably absent from the HIMI dataset, with all samples retaining the avian-like 627E residue. However, other mammalian-adaptive markers were ubiquitous across the HIMI dataset. These include PB1-F2:N66S, associated with increased disease severity in mammals by enhancing inflammation and inhibiting the early immune responses^30^ and the PA mutations S224P and N383D which work together to significantly enhance replication efficacy in mammalian cells^33^. The nucleoprotein (NP) I41V mutation linked to viral replication at lower temperatures of the mammalian upper respiratory tract (as opposed to the higher temperatures typical of the avian intestinal tract) was also present in HIMI viruses^30,34,35^. Finally, all samples shared several HA mutations (HA1 S133A, S154N, T156A and HA2 K64E) that may improve replication in mammals^30,34,35^.

## DISCUSSION

HPAI H5N1 clade 2.3.4.4b has spread rapidly and widely across the globe and has now been detected at the remote territory of Heard Island and McDonald Islands in the southern Indian Ocean. This is the first detection of the panzootic clade 2.3.4.4b HPAI in Australian sub-Antarctic territory. We confirmed the presence of H5N1 in southern elephant seals, Antarctic fur seals, gentoo and king penguins, a brown skua and a South Georgia diving petrel, and observed mass mortality in elephant seals and elevated mortality in king and gentoo penguins. Our results provide strong evidence that the virus was introduced from Crozet Islands (<1,700 km from HIMI) based on current availability of virus sequences. It is clear from our observations and those of others^2,12,36^ that HPAI can and does spread across vast ocean basins. While the transport host(s) cannot be confirmed, southern giant petrels are a plausible candidate and have been suspected as long-distance transport hosts elsewhere^6,12^. As the virus progresses eastwards around the sub-Antarctic, including along circumpolar flyways, there is potential for further geographic spread to additional HPAI-free regions, including Macquarie Island, New Zealand, Australia and East Antarctica.

Although HPAI was detected in six different wildlife species, mass mortality was only observed in elephant seals. The balance of host, pathogen and environmental factors that drive this pattern remains unknown, highlighting the need to better understand the biological and epidemiological determinants of disease severity and population-level impact. Elephant seals have been disproportionately impacted by H5N1 compared to other Southern Ocean species, which may be due to aspects of their biology and life-history. Compared to other pinniped species, they spend most of their time at sea, but congregate in large, dense colonies to breed and moult^37,38^, creating conditions highly conducive to disease transmission through direct contact and environmental contamination. Immunologically naïve pups^39,40^ are expected to be especially susceptible^39,40^. The species’ strong philopatry further compounds this risk as seals are unlikely to abandon infected sites, potentially trapping successive generations in disease hotspots, facilitating sequential annual outbreaks if the virus persists locally^5^. These mortality events are of conservation concern given that the majority of the global breeding southern elephant seal population is concentrated at four geographically separated regions. We also know a single viral introduction can potentially have considerable long-term (∼70 years) population-level consequences in this species as modelled for Peninsula Valdez, a population that was increasing prior to HPAI^20^. It is precisely these concerns that prompted the upgrade of the global conservation status of southern elephant seals from Least Concern to Vulnerable^41^. So far, the effects of HPAI on southern elephant seals have been modelled for increasing populations. The effect of HPAI on a decreasing population, such as Macquarie Island^41^, remain unclear and of considerable concern.

Though impacts of HPAI on wildlife have varied, some common patterns are emerging within sub-Antarctic and Antarctic ecosystems. Our observations are consistent with those of other sites in certain respects: 1) mass mortality has been observed multiple times in southern elephant seals^20,21^; 2) king and gentoo penguins have consistently exhibited patchy patterns of elevated mortality of a much lower magnitude^6,12,23,42^ but still above baseline level for these species^43^; 3) giant petrels and sheathbills have suffered relatively low mortality^16^ despite live individuals testing positive; 4) mortality patterns in other species (e.g. Antarctic fur seal, skuas, black-browed albatross, shags) have been less consistent across sites, with notable mortalities recorded at some sites and limited mortality at others^12,23,42^. It is not clear to what extent the observed mortality patterns reflect species predilection, viral strain fitness or environmental factors such as habitat feature and timing of introduction. The pattern of virus genetic mutations identified in our H5N1 sequences were identical with those at other sub-Antarctic islands^13^ including several mutations associated with enhanced adaptation to mammalian infection^6,8,16^. The PB2 R251K mutation fixed across the dataset and two elephant seal individuals with PA 266L and NS K55 were the only exceptions.

The spatiotemporal patterns of mortality and the viral phylodynamics both suggest a recent introduction to Heard Island (August 2025), shortly before the expedition arrival in October 2025. Initially, high elephant seal mortality appeared spatially restricted to the south-eastern quadrant, though an isolated HPAI-positive pup carcass in north-western Atlas Cove confirmed island-wide presence. Carcass abundance and age in January 2026 indicated that pup mortality occurred across the island in October 2025. The trend of increasing daily mortality rates at sites progressively further from Winston Lagoon, combined with the highest king penguin mortality in south-eastern colonies, supports the hypothesis that this region was the likely origin of introduction. This is consistent with a recent single source of introduction, supported by the phylodynamic model’s narrow estimated tMRCA of July–September 2025. The apparent arrival of HPAI at HIMI roughly a year after other islands in the southern Indian Ocean is notable, particularly given the close phylogenetic relationship to the two strains from samples collected during necropsies on Crozet prior to that island’s main outbreak^13^. Introduction via unsurveyed nearby islands such as Île de l’Est or Île aux Cochons (in the Crozet Islands group) remain possible. However, the HIMI genomes had lower-than-expected root-to-tip divergence compared with other H5N1 2.3.4.4b viruses. This is most consistent with a low-cadence transmission chain involving few hosts, either on land or at sea, though incomplete sampling of intermediate viruses may also contribute. Hotspots of infection likely generate many such chains of transmission as infected animals disperse, though most likely die out without seeding new, distant outbreaks.

Aerial drone surveys were a powerful, non-invasive technique for identifying and monitoring mortality across seabird and seal populations at HIMI. This remote and environmentally sensitive sub-Antarctic location presents significant logistical challenges, making traditional ground-based surveys difficult at scale and during short visits. Drone platforms enabled broad spatial coverage, including areas inaccessible by foot and offshore islands, and repeat surveys allowed temporal progression of the outbreak to be quantified in elephant seals. They also detected spatially dispersed king penguin mortalities among the species’ dense breeding colonies that were undetectable on foot without causing significant disturbance. High-resolution optical and thermal imaging facilitated systematic surveying with minimal disturbance, but thermal signatures alone proved unreliable for identifying mortality. For example, recently hauled-out live animals exhibited low thermal signatures, while sun exposed carcasses could produce elevated thermal signals. We therefore used unpaired pups as a proxy for dead or moribund individuals, though this likely underestimated true mortality as ground counts confirmed some dead pups were still paired with females. Assigning mortality status to seabird chicks was similarly challenging and elevated chick mortality attributable to HPAI could not be formally assessed, especially given high baseline variability in breeding success^44^. Nevertheless, these findings indicate the emerging role of drones for early detection, risk assessment and ecological monitoring of the impacts of infectious disease in remote wildlife communities. When coupled with the ability to concurrently collect population census data (which was achieved for elephant seals, black-browed albatross, king penguins and shags), drone surveys offer a practical means of rapidly quantifying at-risk populations during an outbreak, a capacity that is frequently absent yet essential for evaluating disease impact^45^.

By assembling this comprehensive multimodal study of an outbreak of HPAI H5N1 affecting multiple species, we demonstrate the utility of drone-based data collection coupled with pathogen genome sequencing and phylodynamics to simultaneously characterise the course and impacts of the outbreak. By combining these tools, it has been possible to estimate the time of H5N1 arrival and quantify both the impacts and the size of the at-risk, data depauperate populations. The emergence of HPAI poses a severe and potentially catastrophic threat to the wildlife of the Southern Ocean and sub-Antarctic environments, where large, dense breeding and moulting aggregations of already vulnerable species, such as albatrosses, penguins, elephant seals and fur seals, create ideal conditions for rapid viral transmission. Thus, once HPAI enters a colony, die-offs can be swift and devastating as in the Heard Island elephant seal population. Long-lived, *k*-selected species, like elephant seals and albatross are of particular concern given their low reproductive rates, late maturation and high site fidelity, as population losses cannot be quickly recovered^20,46^. For species with small, geographically restricted breeding populations, a single severe outbreak in a season could trigger functionally irreversible population collapses. Compounding this is the near impossibility of implementing many disease management tools, such as vaccination, culling of infected individuals and biosecurity barriers, given the scale and remoteness of Southern Ocean breeding sites. This leaves conservation managers with few practical intervention options beyond surveillance and analysis to understand outbreak trajectories. The combination of HPAI’s arrival in the sub-Antarctic with the logistical challenges of managing wildlife health in remote regions makes HPAI one of the more troubling emerging threats to Antarctic and sub-Antarctic ecosystems. These challenges underscore the importance of actions to mitigate other threats that can be managed through sustained investment in targeted, well-designed conservation actions to build the resilience of ecosystems and wildlife populations. We show that the combination of drones and molecular tools can be instrumental in increasing our capability to monitor HPAI movement and species-level impacts and support preparedness for future HPAI incursions.

## METHODS

### Ethics and permits

All work was carried out under University of Tasmania Animal Ethics permit 32155. Access and sampling permits included *Environment Protection and Management Ordinance 1987* permit 25/1118, *Environment Protection and Biodiversity Conservation Act 1999* Part 13 permit E2025/0254 and Australian Import Permits numbers 0007054098 and 0010821795. All BVLOS drone flights were conducted in accordance with CASA.BVLOS.0178.

### Study site

Field work was completed on two voyages to Heard Island (53.0818° S, 73.5042° E) between 12–22 October 2025, and from 31 December 2025 to 22 January 2026, and McDonald Island (53.0414° S, 72.5985° E) on 12 January 2026. Planning took into consideration the likely presence of HPAI on Heard Island prior to arrival and landings were not permitted on McDonald Island. A HPAI Preparedness and Response Plan and a Triggered Action Response Plan were in place, outlining biosafety, biosecurity and sampling protocols for the voyages.

### Drone and ground surveys

Ship and land-based drone surveys were completed at Heard Island and part of McDonald Island. All surveys were conducted by an experienced wildlife ecologist pilot and an observer. Drone operations used quadcopters: DJI M400 aircraft were fitted with either a DJI Zenmuse P1 (with 35 mm lens) or DJI H30T camera and a limited number of flights were completed with a DJI Mavic 3 Enterprise that has an integrated sensor. The M400 drone was operated up to ∼6 km from the pilot, which enabled ship-based drone operations when conditions were not suitable to access Heard Island and to survey McDonald Island. Data collected were either video imagery (including thermal) or photographs. Flights with the M400 were typically a minimum of 80 m vertical separation from wildlife and those with the Mavic 3E no less than 35 m vertical separation, enabling 1 cm ground sample distance. Digital photographs were stitched into orthomosaics using Agisoft Metashape Pro (Agisoft LLC, Russia). For ship-based HPAI surveys, drone imagery was streamed to monitors where other wildlife ecologists could review vision in real-time to assist with observations and help identify animals and areas of interest. Larger species could easily be observed using drones, with limited capacity to assess smaller (e.g. terns) or more solitary, cryptic species (e.g. skuas).

Ground-based surveillance for HPAI was carried out across 10 part-days of field work at Heard Island covering three main areas: Atlas Cove to Cape Gazert, Capsize Beach to Paddick Valley, and Stephenson Lagoon to Compton Lagoon (Fig. 1). Ground searches included a combination of targeted searches for sick or dead animals at breeding colonies, as well as dedicated line searches away from colonies. These searches, along with opportunistic observations during field activities, helped locate smaller or more cryptic species. Ground surveys were restricted to sections of the island that were accessible during the short field window (Fig. 1).

### Estimating mortality and count data

We defined mass mortality as rapidly occurring catastrophic demographic events that punctuate baseline mortality levels ^47^ and elevated mortality as mortality above baseline but not deemed a catastrophic event.

Observations and counts were categorised as 1) systematic counts, whereby every animal of the focal species in an area was assessed as alive or dead to enable proportional mortality estimates; or 2) targeted and opportunistic ground surveys, where colonies were approached and observed for dead or sick animals, however the total number of live animals were not recorded. All observations and counts were made by ecologists familiar with sub-Antarctic wildlife.

Systematic counts were completed using drone imagery, except for a subset of gentoo colonies where DSLR imagery was used. Additional ground counts of elephant seal harems were made along a 1.4 km stretch of Winston Lagoon to complement concurrent drone survey^48^. To assess consistency between these methods, standardised major axis (SMA or Model II regression) regression was performed using the ‘smartr’ package in R^48^ (Fig. S1; Table S2). Manual counts were made according to imagery type: in still images (extracted from drone videos or from ground-captured SLR camera) individuals were annotated in DotDotGoose^49^ while those in orthomosaics were annotated using QGIS (Supplementary Data). In both cases, a grid-search pattern was used to improve accuracy and ensure all areas were surveyed (Supplementary Methods). When survey conditions permitted, sufficient imagery was collected to produce high-resolution orthomosaics; however, video transects of the coastline, were used to supplement coverage when drone operations were restricted. While orthomosaics provide accurate abundance estimates^50^, and those from video are likely to carry greater error, the large size of elephant seals assisted their detection in video. Annotations were reviewed by an independent observer to correct false positives, false negatives or misassignment across age-classes.

During targeted and opportunistic ground surveys, field personnel observed individual animals for signs of HPAI, including incoordination, difficulty breathing, lethargy or other abnormal behaviour. The number and location of any sick or dead animals were recorded. About ∼50% of the Antarctic fur seal population (Stephenson Lagoon to Compton Lagoon; see Fig. 1) was also surveyed.

#### Systematic elephant seal counts

Surveys covered all known breeding areas of elephant seals. Individuals were assigned to appropriate age-sex classes: pup, weaned pup, adult female or adult male. For classes other than pups, individuals were also categorised as alive unless they were obviously dead (Table S1). Due to the difficulty of correctly identifying dead pups in drone imagery, pups were assigned either as paired with a female if they were within one adult body length, or unpaired if they were farther than this distance from an adult female (Fig. 2). This buffer was based on the distribution of separation distances between unweaned pups and adult females in elephant seal colonies on Macquarie Island that were unaffected by HPAI (Supplementary Methods). Only the nearest pup was allocated to a female since twins are rarely encountered in this species^51^. Pups of this age have a high reliance on their mother’s milk and are not sufficiently developed to survive independently, therefore it was assumed that any unpaired pups were either dead or moribund. Spatial proximity rules were used to assign pairing manually in video imagery and via a custom python script in orthomosaics (Fig. 2). Pup mortality was calculated for each harem as the total number of unpaired pups divided by the total number of pups. To avoid disproportionately influencing the mortality estimates, smaller harems (<10 total pups; 21 harems) were excluded from mortality estimates. Areas surveyed with helicopter were also excluded from mortality estimates as pups could not be easily or repeatably identified in this imagery type (14 harems). From October 2025 to January 2026, pup mortality was calculated for three sites (Josephine Beach north and south, and Brown Lagoon) and estimated from the number of carcasses counted during a ground survey in January divided by the total number of pups in October (Table S8). The total pup production in 2025 was estimated as the total number of unpaired pups, weaners and adult females^52^ in all areas surveyed by drone, plus the number of females only (based on the assumption that each breeding female produces a pup) for the areas surveyed by helicopter. To assess spatial variability in mortality, the distance of each harem from the highest mortality harem at Winston Lagoon was calculated (Supplementary Methods).

#### Systematic seabird counts

Heard Island surveys covered all known breeding colonies of black-browed albatross, shags, king penguins (excluding Long Beach) and ∼50% and ∼20% of historically reported gentoo penguin and southern giant petrel colonies, respectively (Table S6). Counts were made from either orthomosaics, still images exported from drone video or ground-captured DSLR photographs (Table S6). Polygons were constructed around seabird colonies in QGIS to delineate the survey area (Fig. S3). The total number of dead and alive adults was manually annotated both within the colony and surrounding areas, except for king penguins whereby the total number of live adults at each colony was calculated using machine learning (Supplementary Methods; Fig. S4). Carcasses were categorised according to a confidence estimate of mortality (dead or suspected dead) and level of decomposition (fresh, some decomposition, desiccated carcass or bones). Desiccated carcasses or bones were not included in mortality estimates (Table S4) as they were assumed to relate to previous mortality of unknown cause. Only adults have been included in reported mortality estimates for all seabirds as chicks are often obscured by their parents, harder to see due to size, or lay splayed making mortality estimates difficult (Fig. S5).

### Sample collection and analysis

#### Sample collection

Samples were collected from fresh carcasses following strict biosecurity and biosafety protocols. In areas where fresh specimens were scarce, older carcasses were also sampled to increase sample sizes. Samples were collected in duplicate, with one sample stored in Universal Transport Medium (UTM; Copan Diagnostics, Murrieta, CA USA) and another in DNA/RNA Shield (Zymo Research Operations, Tustin, CA USA). Bird swabs were collected from 1) oropharynx and cloaca, and 2) brain. Seal swabs were collected from 1) oral and nasal cavity, and 2) brain. All samples were securely packaged in accordance with IATA packaging instruction 650 by accredited personnel, transported to the ship and frozen at −70°C within 24 hours of sampling. Fresh scat samples were also collected opportunistically from live birds and placed in DNA/RNA Shield and stored at −20°C as soon as practical. Both swab and scat samples were sent to the Commonwealth Scientific and Industrial Research Organisation (CSIRO) Australian Centre for Disease Preparedness (ACDP) for Influenza A matrix gene and H5 hemagglutinin gene qPCR testing^53^. PCR positive samples were further characterised by hemagglutinin and neuraminidase sequence subtyping and whole virus genome sequencing.

#### Virus RNA extraction and sequencing

Viral RNA was extracted with the MagMAX-96 viral RNA isolation kit (Thermo Fisher Scientific, Waltham, MA USA) from the samples according to manufacturer’s instructions. Samples that tested positive using the influenza A matrix gene qPCR with a cycle threshold <30 were selected for influenza A virus targeted next- generation sequencing (NGS, Supplementary Data)^54,55^. The AIV genome segments were amplified using the SuperScript III one-step RT-PCR system with high fidelity Platinum Taq DNA polymerase (Thermo Fisher Scientific) and universal influenza A virus gene primers [MBTuni-12 [5’-ACGCGTGATCAGCAAAAGCAGG] and MBTuni-13 [5’- ACGCGTGATCAGTAGAAACAAGG]] as previously described^54,55^.

To enable a rapid turnaround of results, a subset of samples was processed using the Oxford Nanopore rapid barcoding kit (SQK-RBK114.24) and sequenced on R10 MinION flow cells (Oxford Nanopore Technologies, Oxford, UK). For all other samples, dual-indexed DNA libraries were prepared using an Illumina Nextera XT DNA Library Preparation kit and were sequenced with 2 x 150 cycle paired ends on Illumina MiSeq or NextSeq NGS platforms (Illumina, San Diego, CA, USA), according to manufacturer’s instructions. Sequence reads from both Illumina and Oxford Nanopore platforms were trimmed for quality and mapped to reference sequences of influenza A using Geneious Prime software (Biomatters, Auckland, NZ). Consensus sequence data were manually curated/annotated.

#### Phylogenetic analyses

A reference H5N1 2.3.4.4b dataset was constructed based on the backbone described by Clessin et al. (2025) with a focus on South America, Antarctica and the sub-Antarctic islands. Publicly available sequences were obtained from GISAID (gisaid.org) and BV-BRC (bv-brc.org) and combined with the 26 newly generated genomes from Heard Island, plus 21 unpublished sequences from Crozet Islands and Kerguelen Islands^13^. A small number of North American high-quality reference sequences were included for broader epidemiological context, while South American, Antarctic and sub-Antarctic Island sequences were retained in full. After quality filtering and removal of duplicates, the final dataset contained 1,322 genomes.

Time-resolved phylogenies were inferred from the concatenated genomes using BEAST X v.10.5.0^56^ following the same overall framework of Clessin et al. (2025) using the same substitution model (GTR+Γ), uncorrelated relaxed molecular clock with an underlying lognormal distribution ^57^, and a Bayesian SkyGrid coalescent prior^58^. Hamiltonian Monte Carlo was used to efficiently infer the SkyGrid^59^ and molecular clock model parameters^60^. Independent Markov chain Monte Carlo (MCMC) analyses of 250 million iterations were run with GPU acceleration using the BEAGLE v.4.0.1^61^ high-performance computational library, and combined after removal of burn-in. A combined number of 1,000 posterior trees were sampled and stored for subsequent phylogeographic analysis. Convergence and proper mixing of independent replicates were assessed in Tracer v.1.7.2^62^ with key parameters including root age, effective population sizes over time and molecular clock model parameters showing satisfactory efficient sample size (ESS) values >200. Convergence in phylogenetic tree space was assessed using multidimensional scaling projections of posterior tree space based on pairwise Robinson-Foulds distances^63^ between sampled trees^64,65^. Majority-rule highest independent posterior subtree reconstruction (HIPSTR) consensus trees were generated using TreeAnnotator X (v.10.5.0)^66^ and visualised in R v.4.3.1^67^ with ggtree v.3.8.2^68^.

#### Phylogeographic analyses

The posterior trees stored during the phylogenetic analysis were used as an empirical tree distribution to estimate spatiotemporal dispersal patterns using an asymmetric forward-in-time (FIT) discrete phylogeographic model with Bayesian stochastic search variable selection^69^. Independent MCMC chains of 20 million states were run with GPU acceleration using the BEAGLE v.4.0.1^61^ high-performance computational library. Convergence and proper mixing were assessed in Tracer v.1.7.2^62^. Location-annotated majority-rule HIPSTR consensus tree were generated using TreeAnnotator X (v.10.5.0) and visualised in R v.4.3.1^67^ with ggtree v.3.8.2^68^.

#### Virus molecular marker analysis

To identify key molecular markers that may inform host adaptation, increased virulence, antiviral resistance and virus fitness, the reference dataset was screened using the FluMut pipeline^70^, which generates outputs mapping detected markers to their biological effects and relevant literature. Transcribed amino acid sequence of each virus gene segment were aligned in MEGA^71^ and cross checked with calls from the FluMut pipeline. Substitution matrices were then generated for all positions using a custom R script where at least one position differed from a reference. Mutations occurring across all HIMI samples and fixed in the populations were recorded.

## ACKNOWLEDGMENTS

Thanks to Simon Payne and Doug Thost for assisting Jarrod Hodgson with drone operations. Field teams in addition to some authors included Dahlia Foo, Rowena Hannaford, Madi McLatchie, Demelza Wall and Jessica Williams. Thanks to Megan Rigby, Ros Watson and Dion Frampton from CSIRO Hobart, Tristan Reid, Frank Wong and Mark Ford from ACDP, John van den Hoff, Dan Wilkins and Patricia Miloslavich at the Australian Antarctic Division, the Tasmania Parks and Wildlife Service and the Australian Government HPAI Preparedness Taskforce for support. Thanks to Frank Wong and John van den Hoff for comments on the manuscript. This work was part of a larger management field campaign led and funded by the Australian Government Department of Climate Change, Energy, the Environment and Water’s Australian Antarctic Division. We thank all those involved in the campaign through planning, logistics, permitting and operations to facilitate and achieve project objectives. Thierry Boulinier acknowledges support from ANR grants ECOPATHS (ANR-25-CE35-0016) and WILDFLU (ANR-25-CE35-0691), and French Polar Institute (IPEV) and CNRS SEE-Life project ECOPATH-1151. Guy Baele acknowledges support from the Research Foundation - Flanders (“Fonds voor Wetenschappelijk Onderzoek - Vlaanderen,” G098321N), from the European Union Horizon 2023 RIA project LEAPS (grant agreement no. 101094685), and from the DURABLE EU4 Health project 02/2023-01/2027, which is co-funded by the European Union (call EU4H-2021-PJ4) under Grant Agreement No. 101102733. Finally, we gratefully acknowledge all data contributors, i.e., the authors and their originating laboratories responsible for obtaining the specimens, and their submitting laboratories for generating the genetic sequence and metadata and sharing via the GISAID Initiative, on which this research is based.

## DATA AVAILABILITY

Sequence data will be made publicly available on Genbank on acceptance and accession numbers provided. Other data will be available via the Australian Antarctic Data Centre (https://data.aad.gov.au/), with a DOI provided on acceptance.

## Author contributions

Study conceptualisation and design: JM, JH, AT, BW, TBu, CM, TBo. Funding and permitting: AT, JM, JH, BW, LW, AP. Field surveys: JH, JM, MW, GM, AP, HA, RA, AL, BV. Sample collection: JM, JH, MW, GM, LW, AP, JT, ML, RA, AL, BV, PV. Data processing: JH, MW, GM, JM, LW, HA, RA, AL, BV, PV. Sequencing: MN, FXB. Phylogenetic and mutation analysis: MN, GBa, AP. Writing of the original draft: JM, JH, CM, MN, TBu, MW, GM, AP. Draft revision and editing: all authors.

## REFERENCES

1 Couty, M., et al. The role of wild birds in the global highly pathogenic avian influenza H5 panzootic, 2020–2023. NPJ Biodiversity 5 (2026).

2 Kuiken, T. et al. Emergence, spread, and impact of high-pathogenicity avian influenza H5 in wild birds and mammals of South America and Antarctica. Conservation Biology 40 (2026).

3 Gamarra-Toledo, V. et al. Highly Pathogenic Avian Influenza (HPAI) strongly impacts wild birds in Peru. Biological Conservation 286 (2023).

4 Leguia, M. et al. Highly pathogenic avian influenza A (H5N1) in marine mammals and seabirds in Peru. Nature Communications 14 (2023).

5 Uhart, M. M. et al. Epidemiological data of an influenza A/H5N1 outbreak in elephant seals in Argentina indicates mammal-to-mammal transmission. Nature Communications 15 (2024).

6 Bennison, A. et al. A case study of highly pathogenic avian influenza (HPAI) H5N1 at Bird Island, South Georgia: the first documented outbreak in the subantarctic region. Bird Study 71, 380–391 (2024).

7 Banyard, A. C. et al. Detection and spread of high pathogenicity avian influenza virus H5N1 in the Antarctic Region. Nature Communications 15 (2024).

8 Falklands Conservation. Falklands Conservation Annual seabird monitoring report 2023/2024. (2024).

9 Aguado, B. et al. Searching for high pathogenicity avian influenza virus in Antarctica. Nature Microbiology 9, 3081–3083 (2024).

10 Bennett-Laso, B. et al. Confirmation of highly pathogenic avian influenza H5N1 in skuas, Antarctica 2024. Frontiers in Veterinary Science 11 (2024).

11 Iervolino, M., et al. The expanding H5N1 avian influenza panzootic causes high mortality of skuas in Antarctica. Scientific Reports 16 (2026).

12 Clessin, A. et al. Circumpolar spread of avian influenza H5N1 to southern Indian Ocean islands. Nature Communications 16 (2025).

13 Clessin, A. et al. Dispersal, adaptation and persistence of H5N1 in the sub-Antarctic and Antarctica. bioRxiv, 2026.2003.2020.713283 (2026).

14 Steinfurth, A., et al. Investigating high pathogenicity avian influenza virus incursions to remote islands: detection of H5N1 on Gough Island in the South Atlantic Ocean. Emerging Microbes and Infections 15 (2026).

15 Waller, S. et al. Avian Influenza Virus Surveillance Across New Zealand and Its Subantarctic Islands Detects H1N9 in Migratory Shorebirds, but Not 2.3.4.4b HPAI H5N1. Influenza and other Respiratory Viruses 19 (2025).

16 SCAR. Sub-Antarctic and Antarctic Highly Pathogenic Avian Influenza H5N1 Monitoring Project, 2026).

17 Ryan, P. G. Update on unusual Emperor Penguin mortality at Astrid colony. (Unpublished report to the South African Competent Authority and to IAATO, 16 November 2024) (2024).

18 Vanstreels, R. E. T. et al. Novel Highly Pathogenic Avian Influenza A (H5N1) Virus, Argentina, 2025. Emerging Infectious Diseases 31 (2025).

19 Youk, S. et al. H5N1 highly pathogenic avian influenza clade 2.3.4.4b in wild and domestic birds: Introductions into the United States and reassortments, December 2021-April 2022. Virology 587, 109860 (2023).

20 Campagna, C. et al. Predicting Population Consequences of an Epidemic of High Pathogenicity Avian Influenza on Southern Elephant Seals. Marine Mammal Science 41 (2025).

21 Bamford, C. C. G. et al. Highly Pathogenic Avian Influenza Viruses (HPAIV) Associated with Major Southern Elephant Seal Decline at South Georgia. Communications Biology 8 (2025).

22 Campagna, C. et al. Catastrophic mortality of southern elephant seals caused by H5N1 avian influenza. Marine Mammal Science 40, 322–325 (2024).

23 DFFE. High Pathogenicity Avian Influenza (H5N1) confirmed on Subantarctic Marion Island. (Department of Forestry, Fisheries and Environment, Republic of South Africa) (2025).

24 Lisovski, S. et al. Unexpected Delayed Incursion of Highly Pathogenic Avian Influenza H5N1 (Clade 2.3.4.4b) Into the Antarctic Region. Influenza and other Respiratory Viruses 18 (2024).

25 Tyndall, A. A., et al. Quantifying the Impact of Avian Influenza on the Northern Gannet Colony of Bass Rock Using Ultra-High-Resolution Drone Imagery and Deep Learning. Drones 8 (2024).

26 Hindell, M. A. et al. Circumpolar habitat use in the southern elephant seal: Implications for foraging success and population trajectories. Ecosphere 7 (2016).

27 Chua, M., Ho, S. Y. W., McMahon, C. R., Jonsen, I. D. & de Bruyn, M. Movements of southern elephant seals (Mirounga leonina) from Davis Base, Antarctica: combining population genetics and tracking data. Polar Biology 45, 1163–1174 (2022).

28 Weimerskirch, H., Mougey, T. & Hindermeyer, X. Foraging and provisioning strategies of black-browed albatrosses in relation to the requirements of the chick: Natural variation and experimental study. Behavioral Ecology 8, 635–643 (1997).

29 Li, J. et al. Three amino acid substitutions in the NS1 protein change the virus replication of H5N1 influenza virus in human cells. Virology 519, 64–73 (2018).

30 Suttie, A. et al. Inventory of molecular markers affecting biological characteristics of avian influenza A viruses. Virus Genes 55, 739–768 (2019).

31 Fusade-Boyer, M., et al. Investigation and impact of mammalian adaptation markers on H5N8 high pathogenicity avian influenza polymerase activity. NPJ Viruses 4 (2026).

32 Kim, D. H., Lee, D. Y., Seo, Y., Song, C. S. & Lee, D. H. Immediate PB2-E627K amino acid substitution after single infection of highly pathogenic avian influenza H5N1 clade 2.3.4.4b in mice. Virology Journal 22 (2025).

33 Song, J., Xu, J., Shi, J., Li, Y. & Chen, H. Synergistic effect of S224P and N383D substitutions in the PA of H5N1 avian influenza virus contributes to mammalian adaptation. Scientific Reports 5 (2015).

34 Wang, W. et al. Glycosylation at 158N of the hemagglutinin protein and receptor binding specificity synergistically affect the antigenicity and immunogenicity of a live attenuated H5N1 A/Vietnam/1203/2004 vaccine virus in ferrets. Journal of Virology 84, 6570–6577 (2010).

35 Gao, Y. et al. Identification of amino acids in HA and PB2 critical for the transmission of H5N1 avian influenza viruses in a mammalian host. PLoS Pathogens 5 (2009).

36 Ross, T. A. et al. AviFluMap: An interactive tool to assess H5N1 avian influenza incursion risk in Australia via migratory birds. Ecological Informatics 93 (2026).

37 Carrick, R. & Ingham, S. E. Ecological studies of the southern elephant seal Mirounga leonina (L), at Macquarie Island and at Heard Island. Mammalia 24, 325–342 (1960).

38 Laws, R. M. The elephant seal (Mirounga leonina Lin) II General, social and reproductive behaviour. Falkland Islands Dependencies Survey Scientific Reports 13 (1956).

39 Marquez, M. E. I., Carlini, A. R., Slobodianik, N. H., De Ferrer, P. A. R. & Godoy, M. F. Immunoglobulin M serum levels in females and pups of southern elephant seal (Mirounga leonina) during the suckling period. Comparative Biochemistry and Physiology - A Molecular and Integrative Physiology 119, 795–799 (1998).

40 Hall, A. J. et al. The immunocompetence handicap hypothesis in two sexually dimorphic pinniped species - Is there a sex difference in immunity during early development? Developmental and Comparative Immunology 27, 629–637 (2003).

41 Hofmeyr, G. J. G. Mirounga leonina. In: The IUCN Red List of Threatened Species (2026).

42 FIG. Avian Influenza Information. (ed Falkland Islands Government Falkland Islands Department of Agriculture) (2026).

43 Le Bohec, C. et al. Population dynamics in a long-lived seabird: I. Impact of breeding activity on survival and breeding probability in unbanded king penguins. Journal of Animal Ecology 76, 1149–1160 (2007).

44 Bardon, G. et al. Multiannual environmental forcing shapes breeding phenology and success in a sub-Antarctic seabird. Science Advances 12, 1–11 (2026).

45 Guberti, V., Stancampiano, L. & Ferrari, N. Surveillance, monitoring and survey of wildlife diseases: A public health and conservation approach. Hystrix 25 (2014).

46 McMahon, C. R., Hindell, M. A., Burton, H. R. & Bester, M. N. Comparison of southern elephant seal populations, and observations of a population on a demographic knife-edge. Marine Ecology Progress Series 288, 273–283 (2005).

47 Fey, S. B. et al. Recent shifts in the occurrence, cause, and magnitude of animal mass mortality events. Proceedings of the National Academy of Sciences 112, 1083–1088 (2015).

48 Warton, D. I., Duursma, R. A., Falster, D. S. & Taskinen, S. smatr 3- an R package for estimation and inference about allometric lines. Methods in Ecology and Evolution 3, 257–259 (2012).

49 DotDotGoose (American Museum of Natural History, Center for Biodiversity and Conservation. Available from http://biodiversityinformatics.amnh.org/open_source/dotdotgoose., version 1.7.0).

50 Hodgson, J. C. et al. Drones count wildlife more accurately and precisely than humans. Methods in Ecology and Evolution 9, 1160–1167 (2018).

51 McMahon, C. R. & Hindell, M. Twinning in southern elephant seals: the implications of resource allocation by mothers. Wildlife Research 30, 35–39 (2003).

52 McCann, T. S. in Studies of sea mammals in south latitudes. Proceedings of a symposium of the 52nd. Vol. May 1982 (eds J.K. Ling & M.M. Bryden) 1–17 (ANZAAS Congress in Sydney, 1985).

53 Heine, H. G., Trinidad, L., Selleck, P. & Lowther, S. in Avian Diseases. SUPPL. 1 edn 370–372.

54 Kampmann, M. L. et al. A simple method for the parallel deep sequencing of full influenza A genomes. Journal of Virological Methods 178, 243–248 (2011).

55 Zhou, B. et al. Single-reaction genomic amplification accelerates sequencing and vaccine production for classical and swine origin human influenza A viruses. Journal of Virology 83, 10309–10313 (2009).

56 Baele, G. et al. BEAST X for Bayesian phylogenetic, phylogeographic and phylodynamic inference. Nature Methods 22, 1653–1656 (2025).

57 Drummond, A. J., Ho, S. Y. W., Phillips, M. J. & Rambaut, A. Relaxed phylogenetics and dating with confidence. Plos Biology 4, 699–710 (2006).

58 Gill, M. S. et al. Improving Bayesian Population Dynamics Inference: A Coalescent-Based Model for Multiple Loci. Molecular Biology and Evolution 30, 713–724 (2013).

59 Suchard, M. A., Baele, G., Gill, M. S. & Lemey, P. Hamiltonian Monte Carlo sampling to estimate past population dynamics using the skygrid coalescent model in a Bayesian phylogenetics framework. Wellcome Open Research 5 (2020).

60 Ji, X. et al. Gradients Do Grow on Trees: A Linear-Time O(N)-Dimensional Gradient for Statistical Phylogenetics. Molecular Biology and Evolution 37, 3047–3060 (2020).

61 Ayres, D. L. et al. BEAGLE 3: Improved Performance, Scaling, and Usability for a High-Performance Computing Library for Statistical Phylogenetics. Systematic Biology 68, 1052–1061 (2019).

62 Rambaut, A., Drummond, A. J., Xie, D., Baele, G. & Suchard, M. A. Posterior summarization in Bayesian phylogenetics using Tracer 1.7. Systematic Biology 67, 901–904 (2018).

63 Robinson, D. F. & Foulds, L. R. Comparison of phylogenetic trees. Mathematical Biosciences 53, 131–147 (1981).

64 Gao, J. et al. Biological causes and impacts of rugged tree landscapes in phylodynamic inference. Proceedings of the National Academy of Sciences 123, e2510938123 (2026).

65 Brusselmans, M. et al. On the importance of assessing topological convergence in Bayesian phylogenetic inference. Virus Evolution 10 (2024).

66 Baele, G. et al. HIPSTR: highest independent posterior subtree reconstruction in TreeAnnotator X. Bioinformatics 41, btaf488 (2025).

67 R: A Language and Environment for Statistical Computing v. https://www.R-project.org/ (R Foundation for Statistical Computing, Vienna, Austria (2024).

68 Yu, G., Smith, D. K., Zhu, H., Guan, Y. & Lam, T. T. Y. ggtree: an r package for visualization and annotation of phylogenetic trees with their covariates and other associated data. Methods in Ecology and Evolution 8, 28–36 (2017).

69 Lemey, P., Rambaut, A., Drummond, A. J. & Suchard, M. A. Bayesian phylogeography finds its roots. PLoS Computational Biology 5 (2009).

70 Kumar, S. et al. MEGA12: Molecular Evolutionary Genetic Analysis Version 12 for Adaptive and Green Computing. Molecular Biology and Evolution 41 (2024).

71 Giussani, E. et al. FluMut: a tool for mutation surveillance in highly pathogenic H5N1 genomes. Virus Evol 11, veaf011 (2025).

