## Supplementary material for "Mass mortality of southern elephant seals during multi-species outbreak of HPAI H5N1 on sub-Antarctic Heard Island"

**SUPPLEMENTARY METHODS**

*Systematic southern elephant seal counts*

Systematic counts of elephant seals within harems were undertaken from video imagery, still images and orthomosaics. Video imagery was captured using three video types: wide, zoomed and thermal. From these video types, still images were extracted for each harem to count individuals. For video and still images DotDotGoose[1] was used to digitise individuals into defined age-sex classes (Table S1), along with individual status (alive or dead). Thermal imagery was used to assist with the identification and differentiation of individuals, but not to inform age class or status (e.g. a heat signature does not equate to an alive individual). Due to their larger size, dead adults were obvious from the aerial imagery by the presence of scavengers and/or the state of decomposition. To maintain conservative estimates, only those with high certainty were classed as dead, all others were assumed alive. Dots were placed in the centre of the individual in both the x and y axis. All counters were trained on the same dataset, using the same technique, and training counts were reviewed and calibrated by the same reviewer. Areas of variability were identified in the training dataset, and definitions and rules were developed to avoid misinterpretation by counters (Table S1). For orthomosaics, QGIS was used to manually digitise individuals and categorise them into broad age-classes that were used as inputs for the custom python script that applied spatial proximity rules. Counts from all imagery types were independently reviewed to correct any false positives, false negatives, misassignment and harem boundaries.

Spatial proximity was used to assign pup and adult female pairings. Neonate pups, which are all black and still suckling their mothers, were classed as paired or unpaired with the assumption that an unpaired pup is already dead or will not survive[2]. Pairing distances were informed by those observed in harems at Macquarie Island Nature Reserve unaffected by HPAI in 2022. A distance threshold equivalent to the body length of an adult female (2.85 m; n = 1,634 females measured from Winston and Compton Lagoon orthomosaics) was selected as a conservative maximum distance for assigning pup–female pairings. Using this threshold at Macquarie Island resulted in a mean (± SD) separation distance of 1.24 ± 0.54 m (n = 1,445 pairings across 10 harems).

To assess the accuracy of drone counts, concurrent ground counts were undertaken at a subset of harems along Winston Lagoon. We used a standardised major axis regression (SMA) because both drone and ground counts were subject to error and neither variable was considered independent. Analyses were conducted using counts collected concurrently within the same harem sections, allowing direct comparisons between survey methods. Separate SMA models were fitted for each age-sex class, including total adult females, pups assumed alive, pup assumed dead and total pups. Agreement between methods was assessed using the estimated slope, intercept, confidence intervals and coefficient of determination ($R^2$) (Fig. S1; Table S2). From these models we saw strong correlation between ground and drone counts, except for live pups which were overestimated in drone imagery, however the total pup count aligned between the methods. This was due to some dead pups identified in ground surveys still being paired with a female. As such our mortality estimates are conservative.

**Table S1: Categories used to assign demographic classes to southern elephant seals during systematics counts in DotDotGoose and QGIS from drone imagery collected during October 2025 at Heard Island.**

| Code | Definition (must meet all criteria) | Relevant assumptions | Interpretation of class for population monitoring and reporting | Breeding unit |
|---|---|---|---|---|
| 01a_Pup_paired | a) black in colour +<br>b) paired with an AdF that is located within 1 AdF body length | 1, 2, 4 | A pup that is assumed to be alive and is not weaned (i.e. it is still being fed by its mother). | No |
| 01b_Pup_unpaired | a) black in colour +<br>b) with no AdF<br>**OR** c) obviously dead | 1, 2, 4 | A pup prior to being weaned that does not have a mother in their vicinity and so is assumed to be dead, dying or soon to be dead. | Yes |
| 02a_Weaner_alive | a) >50% silver in colour +<br>b) with no AdF | 1, 2, 4 | Weaned pup that is assumed to be alive. | Yes |
| 02b_Weaner_dead | a) >50% silver in colour +<br>b) with no AdF +<br>c) obviously dead | 1, 2, 4 | Weaned pup that is dead. | Yes |
| 03a_AdF_alive | | 3, 4 | An adult female that is assumed to be alive. | Yes |
| 03b_AdF_dead | | 3, 4 | An adult female that is dead. | Yes |
| 04a_AdM-Within_alive | a) within 2 x AdF body lengths of an AdF | 3, 4 | A dominant bull that is part/integral to a harem and that is assumed to be alive. | No |
| 04b_AdM-Within_dead | a) within 2 x AdF body lengths of an AdF<br>b) obviously dead | 3, 4 | A dead dominant bull that was part/integral to a harem. | No |
| 05a_M-Adjacent_alive | a) > 2 x AdF body lengths of an AdF<br>b) < 50 m from an AdF | 3, 4 | A male that is close/near to a harem and assumed to be alive and interacting with that harem. | No |
| 05b_M-Adjacent_dead | a) > 2 x AdF body lengths of an AdF<br>b) < 50 m from an AdF<br>c) obviously dead | 3, 4 | A dead male that is close/near to a harem and assumed to have previously interacted with that harem. | No |
| 06a_M-Outside_alive | a) > 50 m from an AdF | 3, 4 | A male not associated to a harem and that is assumed to be alive. | No |
| 06b_M-Outside_dead | a) > 50 m from an AdF<br>b) obviously dead | 3, 4 | A dead male not associated to a harem. | No |
| 07a_Unk_alive | a) does not satisfy criteria for another category | 3, 4 | | No |
| 07b_Unk_dead | a) does not satisfy criteria for another category<br>b) obviously dead | 3, 4 | | No |
| **Assumptions** | 1. AdF have a maximum of 1 pup per season (e.g. no twins).<br>2. Abandoned black pups will die (i.e. 01b are not necessarily dead at the time they were surveyed).<br>3. AdM, M, AdF and weaners are alive unless obviously dead.<br>4. An individual can only be assigned to one class. | | | |

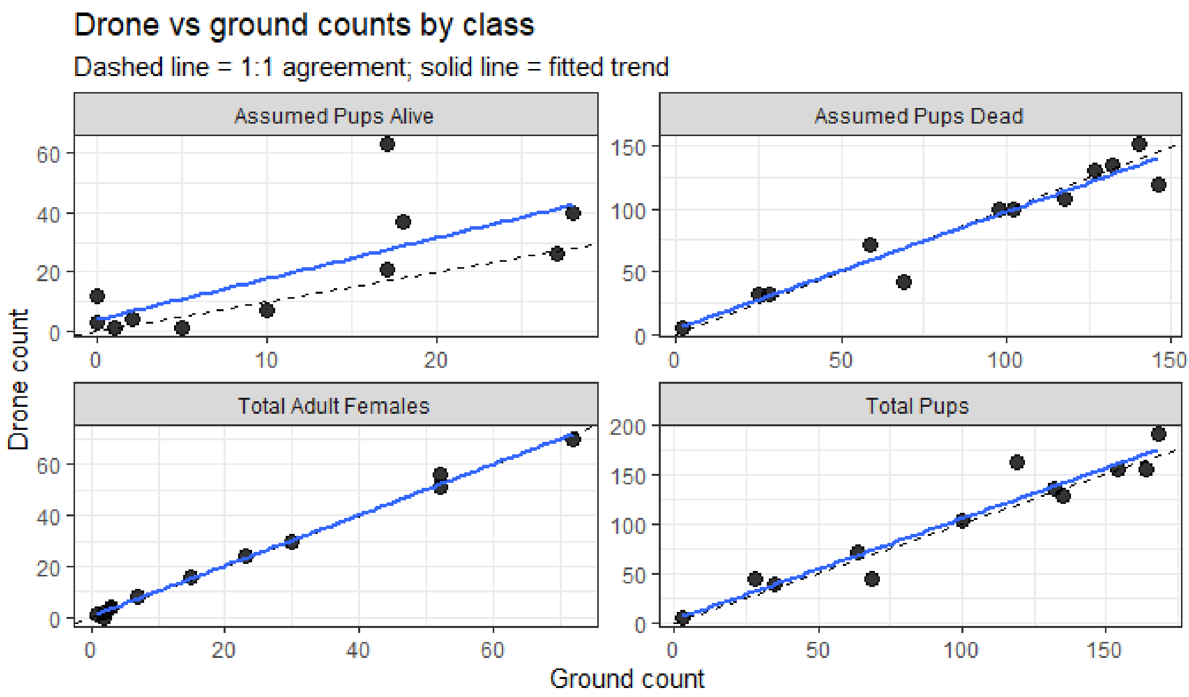

**Fig. S1: Standardized Major Axis (SMA) regression output comparing concurrent drone and ground counts across age-sex classes of southern elephant seals at Heard Island during October 2025.** During ground counts pups were classified as dead or alive, analysis of drone imagery used unpaired and paired pups as an equivalent classification.

**Table S2: Results of standardised major axis (SMA) regressions comparing concurrent drone and ground-based counts across age-sex classes of southern elephant seals at Heard Island during October 2025.** Slopes nearing 1 indicate strong agreement between survey methods.

| Category | Slope | 95% CI (Slope) | Elevation | $R^2$ | p-value |
|---|---|---|---|---|---|
| Assumed Pups Alive | 1.90 | 1.16–3.13 | -2.08 | 0.529 | 0.011 |
| Assumed Pups Dead | 0.97 | 0.80–1.16 | 1.19 | 0.930 | <0.001 |
| Total Adult Females | 1.00 | 0.95–1.05 | 0.38 | 0.996 | <0.001 |
| Total Pups | 1.06 | 0.88–1.29 | -0.66 | 0.922 | <0.001 |

*Cumulative distance calculation*

To investigate spatial patterns in pup mortality across Heard Island, cumulative distance was calculated from the harem with the highest observed pup mortality (SESE_OM14, at Winston Lagoon) using the shortest path. Two methods of calculating the shortest path were assessed, 1) the shortest path around the coastline, reflective of elephant seal distribution, and 2) the least-cost path, representative of a bird transiting between harems (Fig. S2). The first method involved calculating the cumulative distance between harems, originating at SESE_OM14 and using an anticlockwise sequence except for the few harems on the same beach which were to the south-west. The least-cost path was calculated from the flight-path distance from SESE_OM14 to each other harem, constrained to a corridor polygon representing suitable movement habitat bounded by a 1.5km coastal buffer (seaward side) and the 200m elevation contour. The corridor polygon was rasterised at a fixed resolution (10m), with passable cells identified and all others treated as barriers. The shortest-path distances were calculated using the Theta algorithm[3]. Unlike conventional grid-based pathfinding, Theta* incorporates line-of-sight checks during path construction to produce straight-line segments wherever the corridor is unobstructed, avoiding the artificial staircase artefacts that arise from fixed grid connectivity. A post-hoc shortcutting pass was subsequently applied to each path to remove any residual redundant waypoints, with path length recomputed from the final path geometry. All path segments were validated against the rasterised corridor to ensure no segment crossed impassable terrain.

To assess potential spatial patterns in pup mortality, Generalised Additive Models (GAMs) were fitted in R using the 'mgcv' package[4]. Models used a quasibinomial error distribution, with pup mortality modelled as the proportion of unpaired pups relative to total observed pups, using paired and unpaired counts as the binomial response (Table S3).

Separate spatial models were fitted for the shortest path around the coastline, and the least cost path cumulative distances between each harem and SESE_OM14. Smooth terms were fitted with a maximum basis dimension of k = 5, and models were fitted using restricted maximum likelihood (REML).

Although both cumulative distance metrics were significantly associated with pup mortality, the shortest-path distance around the coastline provided the strongest model fit, explaining the greatest proportion of deviance (Table S3). Consequently, this metric was retained as the cumulative distance measure for subsequent analyses and visualisation.

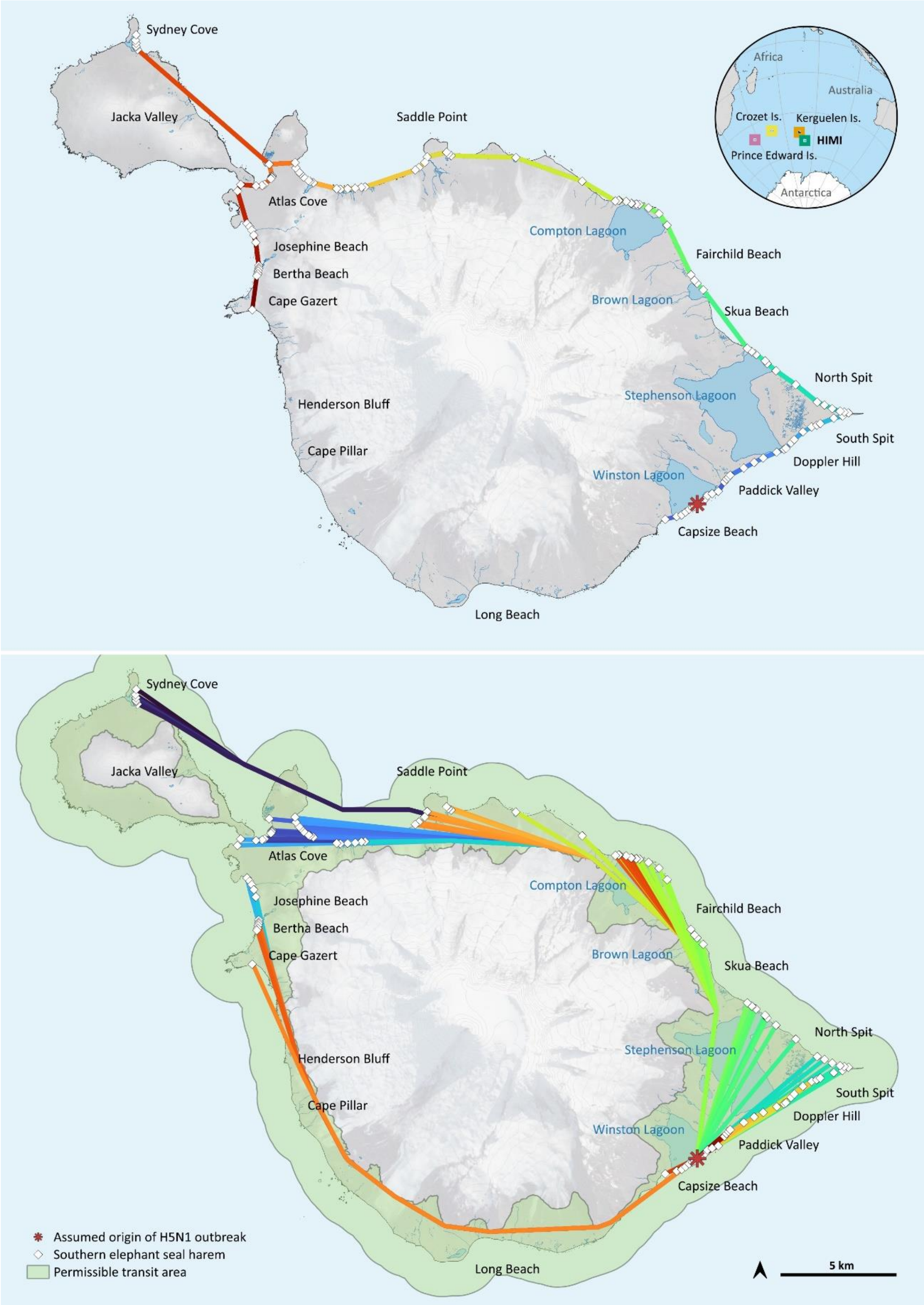

**Fig. S2: Cumulative distances from the harem with the highest mortality (indicated by red star), representative of possible seal-to-seal transmission via each harem, or the shortest path following the coast (top) and transmission via birds, with the least-cost route 'as the bird flys' between each harem (bottom).**

**Table S3: Output from generalized additive models exploring the spatial relationships of the two methods of possible transmission to the proportion of unpaired pups to total pups.** The shortest path around the coastline is a proxy for seal-to-seal transmission via the harems and the least-cost path is a direct line of sight, 'as the bird flys' representing possible transmission via flying birds.

| Model | n | edf | F | p-value | Adjusted R² | Deviance explained | REML | Scale estimate |
|---|---|---|---|---|---|---|---|---|
| Shortest path around the coastline | 74 | 3.64 | 87.72 | <0.001 | 0.950 | 92.6% | -50.62 | 6.60 |
| Least-cost path | 74 | 3.45 | 43.24 | <0.001 | 0.856 | 84.6% | -24.80 | 12.21 |

*Systematic seabird counts*

Orthomosaics were used to count colonies of southern giant petrels (n = 5; all surveyed in January), king penguins (n = 4 in October; n = 8 in January 2026), gentoo penguins (n = 6 in January 2026) and Heard Island shags (n = 4 in October; n = 6 in January 2026). In QGIS, a grid (8 x 8$m^2$) was overlaid on the orthomosaics to encompass the colony and adjacent areas with bird aggregations and each cell was searched (Fig. S3). Still imagery obtained from drones was used to count black-browed albatross colonies (n = 4 in October; n = 3 in January 2026), except for Henderson Bluff where both still imagery and an orthomosaic were utilised. Still imagery obtained from a DSLR camera, taken from a vantage point during ground surveys and stitched in Adobe Photoshop, was used to count additional gentoo colonies not covered by orthomosaics (n = 6 in January 2026). Still images were counted using a grid overlay in DotDotGoose.

For southern giant petrels and black-browed albatross, birds were categorised into age and breeding classes: adult, chick, bird on nest, bird with chick and partner at nest. Due to the difficulty in identifying individual nesting status for shags and gentoo penguins at the time the surveys were conducted, individuals were classed as alive (chick or adult) or dead.

To estimate the total number of live adult king penguins, we used a "You Only Look Once" detection framework model in YOLOv11[5]. Model training and predictions were completed using a custom Python-based package in QGIS (Esparon, Hodgson & Koh, *in prep*). The model was trained using drone imagery from the Doppler Hill colony (Fig. S4) and annotations were predicted using an initial model and then manually corrected (i.e. false positives removed, false negatives corrected). The training and validation datasets included 19,068 and 16,506 annotations, respectively. The resulting model was used to estimate the number of live individuals within the main colony area as well as surrounding areas where congregations of birds were present (Hodgson et al., *in prep*), for all colonies surveyed in January 2026 (Heard Island, n = 8; McDonald Island, n = 1). Consequently, estimates represent the approximate total number of adult individuals, rather than breeding birds only. A confidence threshold of 0.25 was applied to model predications, with detections below this threshold excluded. Predictions were manually reviewed, and obvious false positives were removed.

Each suspected dead individual was assigned a number based on its level of confidence as to whether it was definitely dead (1) or suspected dead (2), decomposition (1-3) and age (adult, chick or unknown) (Table S4). Dead individuals were only categorised into species or age classes if the carcass was distinguishable (i.e. decomposition = 1 or 2). For king penguins, each grid cell was resurveyed by a reviewer to identify any missed individuals and to assess misassignment. Any discrepancies were resolved by a third reviewer. For gentoo penguins all identified dead individuals were reassessed by a reviewer.

To determine whether the observed adult king penguin mortality exceeded baseline levels, data from Macquarie Island was used as a reference. Using orthomosaics constructed from drone imagery of the Lusitania Bay (54.7163° S, 158.8505° E) king penguin colony we undertook a systematic search (area = 10.45 ha) following the same methodology. We assessed two adjacent colonies that were surveyed on 4 January and 2 February 2024.

Only adult mortality has been reported in the manuscript due to the common chick behaviour of laying splayed and seemingly lifeless (Fig. S5), presumably as a means of thermoregulation or energy efficiency. Although some partially decomposed chick carcasses could be identified by the presence of down, due to the uncertainties between distinguishing the splayed live individuals and freshly deceased chicks, they were excluded from further analysis. Only adults have been included in live and dead estimates, count data for each species and population are provided (Tables S5; Table S6).

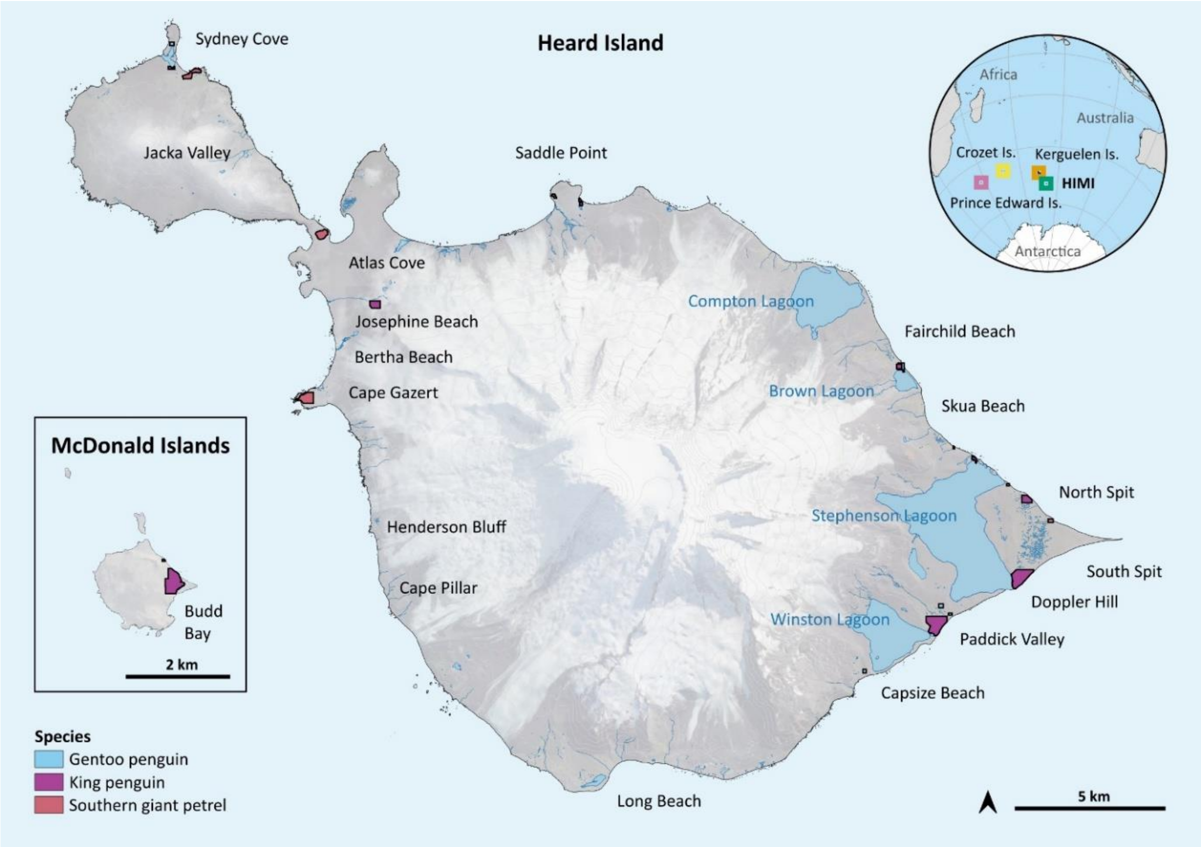

**Fig. S3: Polygons delineating systematic search boundaries used to scan populations of gentoo penguin (n = 6), king penguin (n = 9) and southern giant petrel (n = 5) at Heard Island and McDonald Islands to inform mortality estimates**

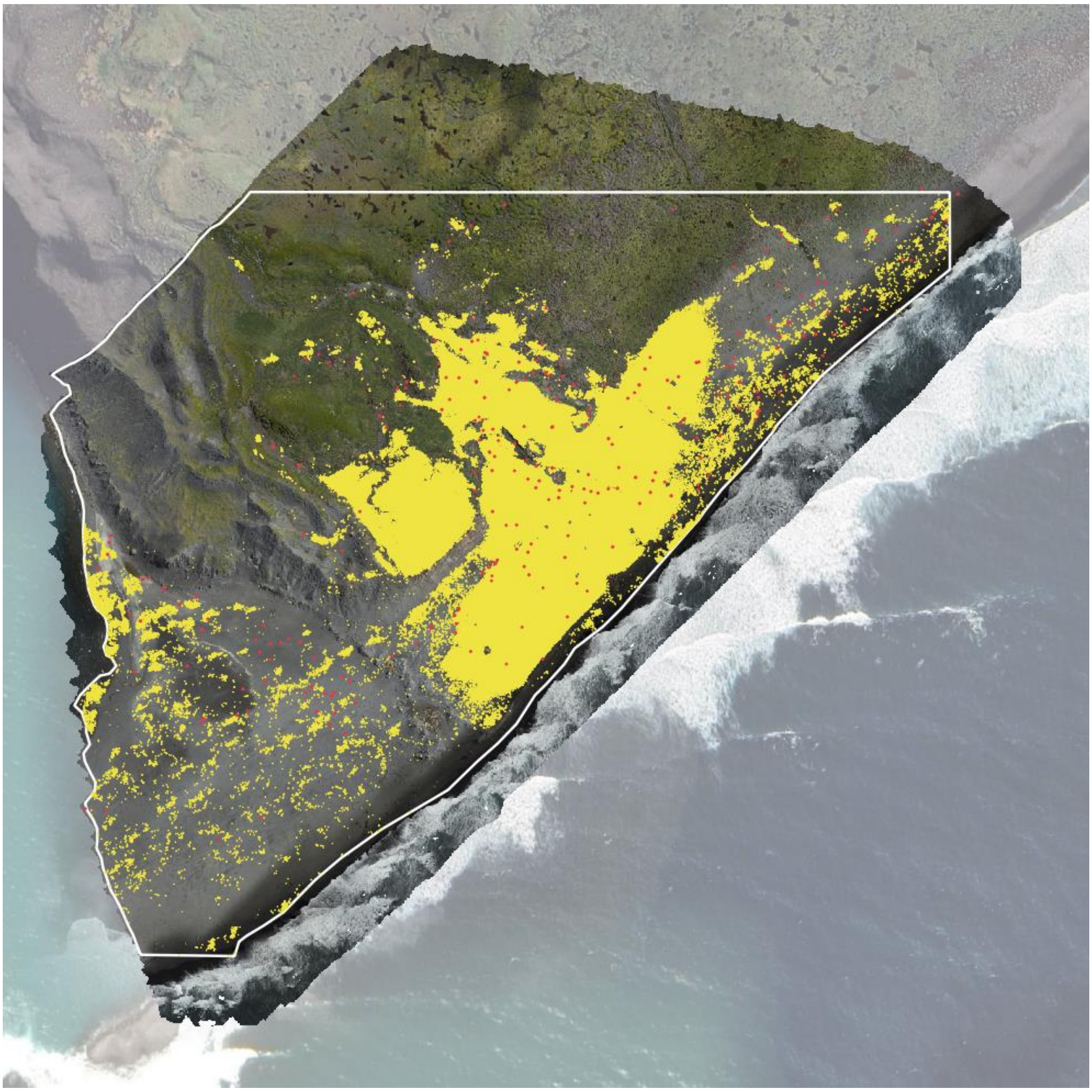

**Fig. S4: Visualisation of the Doppler Hill King penguin colony at Heard Island, generated from the YOLOv11 detection model, used to estimate the total number of alive adults.** The search area boundary (white outline), alive adults predicted by the model (yellow dots; n = 122,271) and manually detected dead adults (red dots; n =253) are shown

**Table S4: Categories used to classify dead seabirds on Heard Island and McDonald Islands during systematic surveys in QGIS.**

| Field | Category | Explanation |
|---|---|---|
| Certainty of mortality | 1. Dead<br>2. Suspected dead | 1. Individuals were classified as dead when they showed visible indicators consistent with death, including loss of normal posture, collapse, lack of body integrity or other indicators inconsistent with a live, resting individual.<br>2. Individuals were classified as suspected dead when they exhibited partial or ambiguous indictors suggesting mortality, but could not be reliably distinguished from live, resting individuals based on the available imagery. |
| Decomposition | 1. Fresh carcass<br>2. Mildly decomposed<br>3. Highly decomposed | 1. Carcass intact with normal body outline and structure, with no visible signs of decomposition.<br>2. Carcass showing visible signs of decomposition, including partial loss of body integrity and/or discoloration, however species and/or age still distinguishable.<br>3. Carcass in an advanced state of decomposition, with substantial loss of body integrity and significant exposure of skeleton. Cannot distinguish age or species with certainty. NB: Individuals in this category were not included in mortality estimates. |

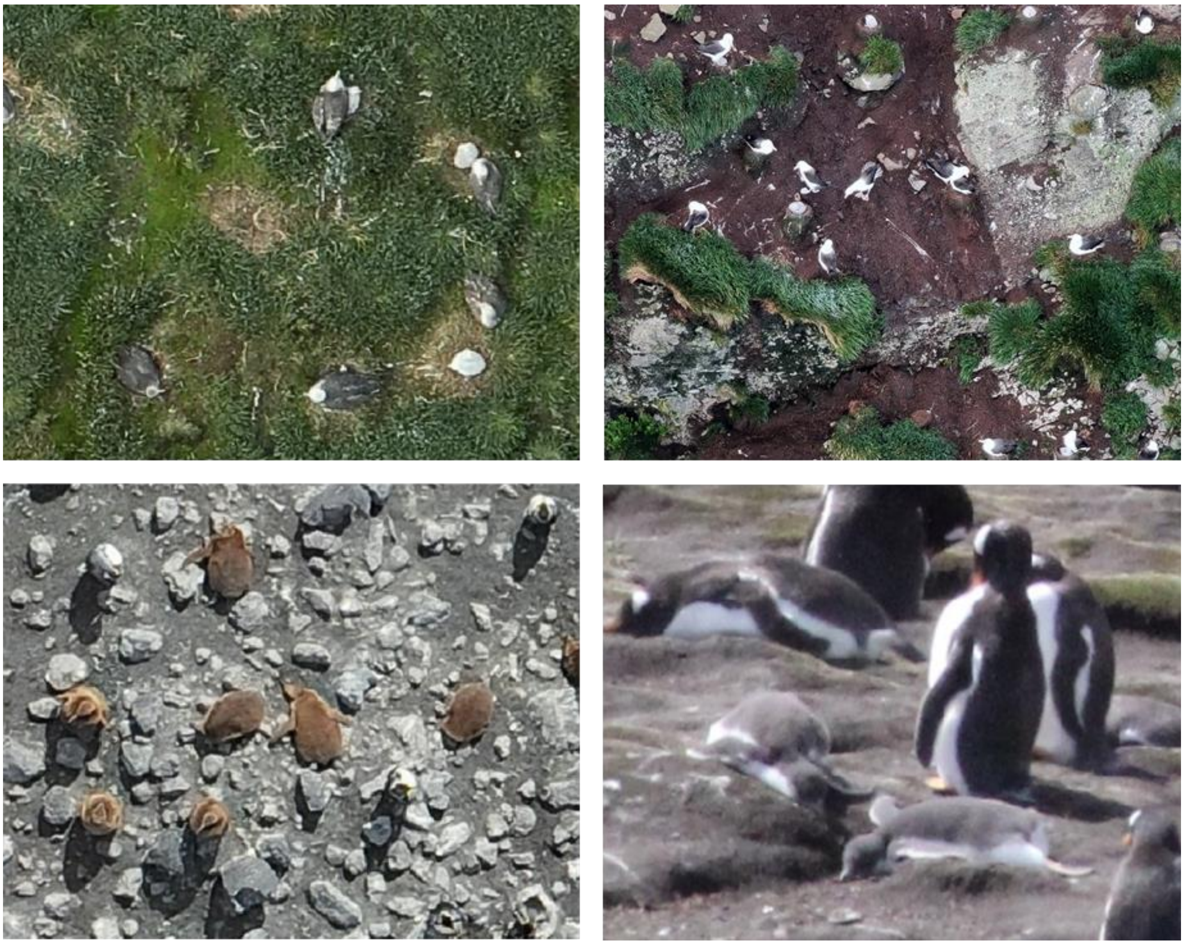

**Fig. S5: Imagery from southern giant petrels (top left), black-browed albatross (top right), king penguins (bottom left) and gentoo penguins (bottom right) showing difficulty in distinguishing chicks for mortality estimates.** Chicks are either too small to see, partially obscured by brooding parents or laying splayed and seemingly lifeless.

*Maximum-likelihood phylogenies*

Sequences were aligned using MAFFT v.7.490[6] and maximum-likelihood phylogenies were inferred for both individual segments and concatenated whole genomes using IQ-TREE v.2.2.0.5[7] with automated model selection using ModelFinder and 1,000 ultrafast bootstraps (UFBOOT). For the concatenated ML tree, ModelFinder selected the GTR+F+I+G4 model according to the Bayesian Information Criterion (BIC). The same concatenated dataset was used for temporal signal assessment using TempEst v.1.5.3[8], which showed strong clock-like behaviour ($R^2$ = 0.84). To assess potential reassortment, the placement of Antarctic and sub-Antarctic genomes was compared across individual segment trees. No well-supported topological incongruence among island-level clusters was observed.

### SUPPLEMENTARY RESULTS

**Table S5: A summary of deceased animals detected at Heard Island and McDonald Islands in October 2025 and January 2026.** Data are either systematic counts of imagery whereby all individuals of the target species were counted within a defined area (i.e., colony and periphery); or counts of dead animals during targeted and opportunistic ground surveys. These data represent total adults (for seabirds) or pups (for southern elephant seals), but do not represent a breeding population estimate.

| **Systematic counts from drone surveys** | | | | |
|---|---|---|---|---|
| | **October 2025** | | **January 2026** | |
| **Species** | **Total live adults** | **Total dead adults** | **Total live adults** | **Total dead adults** |
| Heard Island shag | 1703 | 0 | 1007 | 0 |
| King penguin | | 5 | 235,123 | 298[#] |
| Black-browed albatross | 1256 | 1 | 1426 | 0 |
| Southern giant petrel | | | 1749 | 1 |
| Gentoo penguin* | | | 5125 | 21 |
| **Species** | **Total paired pups^** | **Total unpaired pups^** | | |
| Southern elephant seal | 5,310 | 8,573 | | |
| **Targeted and opportunistic ground surveys** | | | | |
| | **October 2025** | | **January 2026** | |
| **Species** | | **Total dead adults** | | **Total dead adults** |
| Antarctic fur seal | | | | 6 |
| Brown skua | | 1 | | 2 |
| Macaroni penguin | | | | 3 |
| South Georgia diving petrel | | | | 1 |
| Gentoo penguin | | | | 3 |
| King penguin | | 2 | | 16 |
| Southern giant petrel | | 1 | | 2 |

**A subset of gentoo colonies was systematically surveyed using DSLR imagery (n = 6; 2424 alive adults, 2 dead of reported total).*

*^ For the harems where repeated surveys were conducted (n = 16), count data are reported from the most recent timepoint.*

*[#] Includes 18 birds from McDonald Island*

**Table S6: A summary of alive and dead individuals detected at Heard Island and McDonald Islands in October 2025 and January 2026.** Data was obtained using systematic counts of imagery in QGIS or DotDotGoose. Different imagery types are denoted with a footnote, these are stills from a camera (*), stills from drones (#) or orthomosiacs from drones (▪). These data represent total adults (for seabirds) or pups (for southern elephant seals), but do not represent a breeding population estimate.

| Species | Location | October 2025 | | | January 2026 | | |
|---|---|---|---|---|---|---|---|
| | | Survey Date | Live adults | Dead adults | Survey Date | Live adults | Dead adults |
| Heard Island shag | Cape Gazert | | | | 04/01/2026[▪] | 22 | 0 |
| | Cape Pillar | 20/10/2025[#▪] | 1068 | 0 | | | |
| | Gilchrist Beach | 14/10/2025[#] | 65 | 0 | | | |
| | North Spit | 13/10/2025[#] | 298 | 0 | 14/01/2026[▪] | 341 | 0 |
| | Red Island | | | | 10/01/2026[▪] | 27 | 0 |
| | Saddle Point | | | | 14/01/2026[▪] | 73 | 0 |
| | Skua Beach | | | | 15/01/2026[▪] | 344 | 0 |
| | Sydney Cove | 12/10/2025[▪] | 272 | 0 | 10/01/2026[▪] | 200 | 0 |
| | **TOTAL** | | **1703** | **0** | | **1007** | **0** |
| King penguin | Brown Lagoon | | | | 22/01/2026[▪] | 1034 | 0 |
| | Doppler Hill | 16/10/2025[▪] | | 5 | 04/01/2026[▪] | 122271 | 253 |
| | McDonald Island | | | | 12/01/2026[▪] | 32138 | 18 |
| | North Spit | 13/10/2025[▪] | | 0 | 14/01/2026[▪] | 10648 | 5 |
| | Paddick Valley | 21/10/2025[▪] | | 0 | 02/01/2026[▪] | 28805 | 14 |
| | Saddle Point | | | | 14/01/2026[▪] | 2623 | 0 |
| | Schmidt Glacier | 16/10/2025[▪] | | 0 | 04/01/2026[▪] | 33060 | 3 |
| | Stephenson Lagoon | | | | 04/01/2026[▪] | 2358 | 3 |
| | Sydney Cove | | | | 10/01/2026[▪] | 2186 | 2 |
| | **TOTAL** | | | **5** | | 235123 | **298** |
| Black-browed albatross | Charles Carroll Bluff | 12/10/2025[#] | 170 | 0 | 10/01/2026[#] | 225 | 0 |
| | Henderson Bluff | 20/10/2025[▪] | 640 | 1 | | | |
| | Jacka Valley | 19/10/2025[#] | 419 | 0 | 01/01/2026[#] | 379 | 0 |

| Species | Location | October 2025 | | | January 2026 | | |
|---|---|---|---|---|---|---|---|
| | | Survey Date | Live adults | Dead adults | Survey Date | Live adults | Dead adults |
| | | | | | 10/01/2026# | 445 | 0 |
| | | | | | 21/01/2026# | 332 | 0 |
| | Vanhöffen Bluff | 19/10/2025# | 27 | 0 | 10/01/2026# | 45 | 0 |
| | **TOTAL** | | **1256** | **1** | | **1426** | 0 |
| Southern giant petrel | Cape Gazert | | | | 04/01/2026▪ | 1031 | 1 |
| | Mt Aubert De La Rue | | | | 05/01/2026▪ | 152 | 0 |
| | North Spit | | | | 14/01/2026▪ | 156 | 0 |
| | Saddle Point | | | | 14/01/2026▪ | 78 | 0 |
| | Sydney Cove | | | | 10/01/2026▪ | 332 | 0 |
| | **TOTAL** | | | | | **1749** | **1** |
| Gentoo penguin | Brown Lagoon 1 | | | | 15/01/2026* | 390 | 1 |
| | Brown Lagoon 2 | | | | 22/01/2026▪ | 714 | 13 |
| | Capsize Beach | | | | 1/02/2026▪ | 499 | 3 |
| | Fairchild Beach 1 | | | | 15/01/2026* | 817 | 0 |
| | Fairchild Beach 2 | | | | 15/01/2026* | 369 | 0 |
| | Fairchild Beach 3 | | | | 15/01/2026* | 552 | 0 |
| | Gauss Beach | | | | 1/04/2026* | 274 | 1 |
| | North Spit 1 | | | | 14/01/2026▪ | 154 | 0 |
| | Paddick Valley 2 | | | | 1/02/2026* | 22 | 0 |
| | Paddick Valley 3 | | | | 1/02/2026▪ | 254 | 0 |
| | Paddick Valley 4 | | | | 1/02/2026▪ | 518 | 1 |
| | Sydney Cove 1 | | | | 1/10/2026▪ | 555 | 2 |
| | **TOTAL** | | | | | **4404** | **21** |

| Species | Location | October 2025 | | | January 2026 |
|---|---|---|---|---|---|
| | | Survey Date | Paired Pups | Unpaired Pups | |
| Southern elephant seal | Atlas Cove | 10/12/2025# | 268 | 3 | |
| | Capsize Beach | 21/10/2025■ | 153 | 580 | |
| | Corinthian Bay | 10/12/2025# | 619 | 7 | |
| | Fairchild Beach | 13/10/2025# | 441 | 18 | |
| | Gauss Beach | 10/12/2025# | 114 | 4 | |
| | Gilchrist Beach | 14/10/2025#■ | 847 | 74 | |
| | Hoeseason Beach | 10/12/2025# | 134 | 3 | |
| | Jospehine Beach | 10/12/2025# | 288 | 18 | |
| | North Spit | 13/10/2025# | 2167 | 186 | |
| | | 16/10/2025# | 1962 | 847 | |
| | | 22/10/2025# | 815 | 1421 | |
| | Paddick Valley | 21/10/2025■ | 66 | 191 | |
| | Sydney Cove | 10/12/2025# | 57 | 1 | |
| | Skua Beach | 13/10/2025# | 209 | 48 | |
| | South Spit | 16/10/2025# | 2067 | 2424 | |
| | | 21/10/2025-22/10/2025# | 883 | 4174 | |
| | Stephenson Lagoon | 21/10/2025# | 10 | 12 | |
| | Winston Lagoon | 21/10/2025■ | 406 | 2019 | |
| | | **First counts** | **7846** | **5588** | |
| | | **Final counts^** | **5310** | **8573** | |

** Still imagery captured with a DSLR from a ground vantage point*

*# Still imagery captured with drones*

■ *Orthomosaic*

*^ For those harems that were surveyed across multiple timepoints (n = 16), this total is from the last survey conducted.*

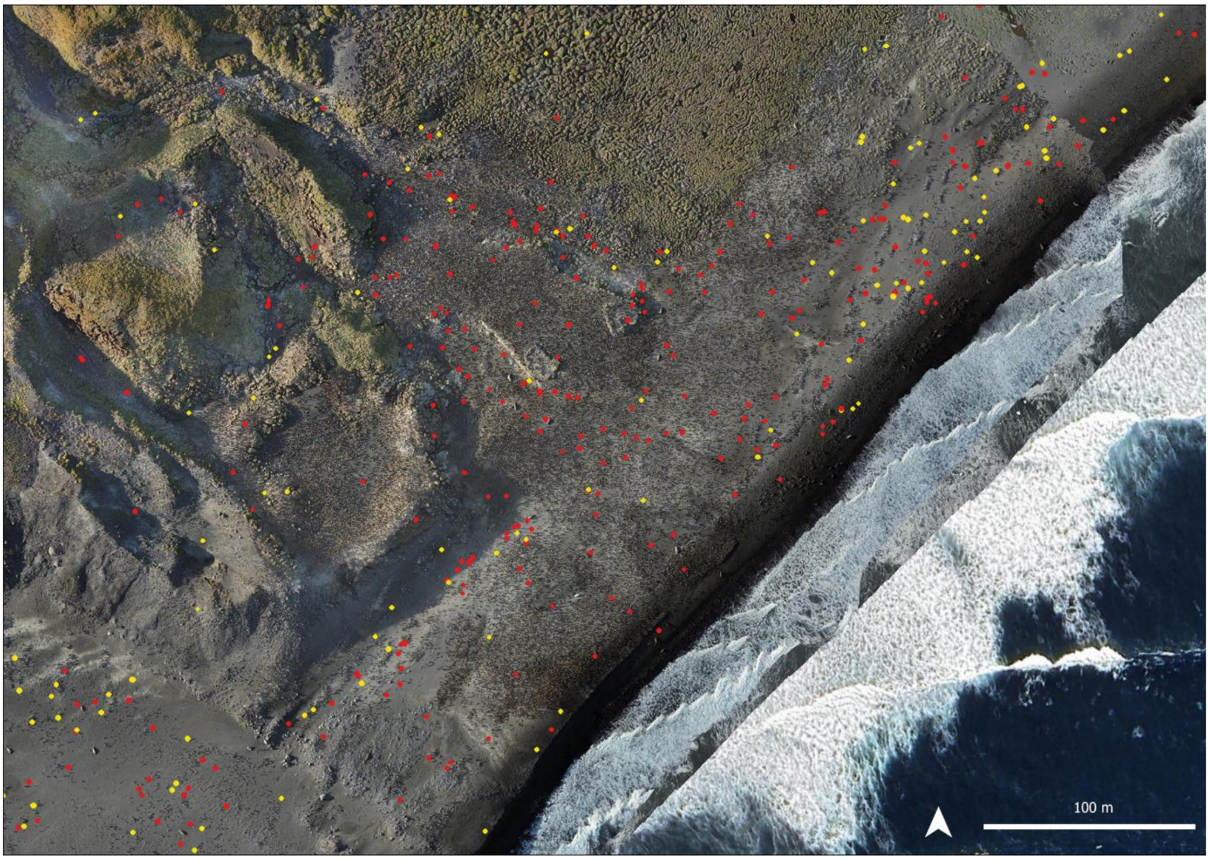

**Fig. S6: Distribution of king penguin carcasses at Doppler Hill, Heard Island in January 2026**. Recently deceased carcasses (decomposition score of 1 or 2) are shown with red dots and old, desiccated carcasses (which were not reported in mortality estimates) are shown with yellow dots.

**Table S7: Breakdown of king penguin systematic mortality counts in October 2025 and January 2026 at Heard Island and McDonald Islands, with both levels of certainty and total area search (hectares) for each colony.**

| | October 2025 | | | January 2026 | | | |
|---|---|---|---|---|---|---|---|
| **Colony** | **Dead** | **Suspected dead** | **Total** | **Dead** | **Suspected dead** | **Total** | **Systematic search area (ha)** |
| Brown Lagoon | | | | 0 | 0 | 0 | 2.58 |
| Doppler Hill | 2 | 3 | 5 | 219 | 33 | 253 | 27.82 |
| McDonald Island | | | | 18 | 0 | 18 | 11.96 |
| North Spit | 0 | 0 | 0 | 4 | 1 | 5 | 7.13 |
| Paddick Valley | 0 | 0 | 0 | 10 | 2 | 14 | 30.79 |
| Saddle Point | 0 | 0 | 0 | 0 | 0 | 0 | 1.85 |
| Schmidt Glacier | 0 | 0 | 0 | 3 | 0 | 3 | 8.02 |
| Stephenson Lagoon | | | | 3 | 0 | 3 | 2.43 |
| Sydney Cove | 0 | 0 | 0 | 1 | 1 | 2 | 1.61 |
| **Total** | **2** | **3** | **5** | **257** | **37** | **298** | **94.20** |

**Table S8: Mortality rates of southern elephant seal pups at sites counted during both October 2025 and January 2026.** October mortality rate was calculated using the ratio of dead pups to total pups. For January estimates, the total breeding units was calculated as the sum of females and dead pups in October, divided by 0.96 to reflect that approximately 96% of pups were assumed to have been born at this stage[9].

| Area | October 2025 | October mortality rate | January 2026 | January mortality rate |
|---|---|---|---|---|
| Josephine Beach Nth | 291 pups (17 dead)<br><br>312 females = 329 units | 5.8% | 211 pup carcasses | 61.6% |
| Josephine Beach Sth | 207 females (~11 dead if similar to Josephine Nth) total 218 units | NA | 194 pup carcasses | 85.4% |
| Brown Lagoon | 101 pups (5 dead)<br><br>116 females | 5% | 99 pup carcasses | 78.5% |

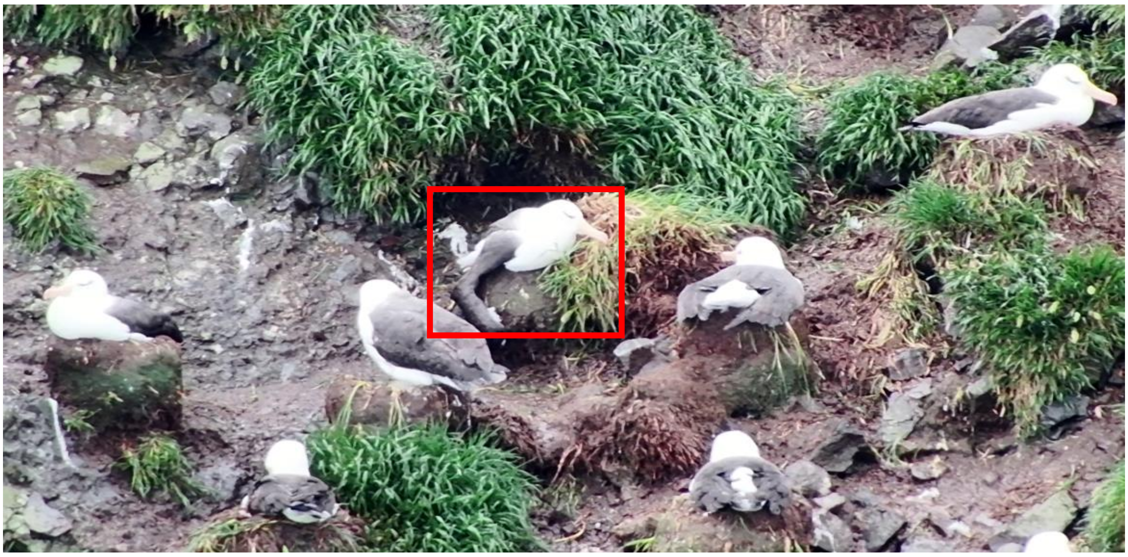

**Fig. S7: Black-browed albatross with a droopy wing on 1 January 2026 at the Jacka Valley colony, Heard Island.** This individual was not on a nest, and no carcass or subsequent unusual behaviour was detected from individuals in the same area during repeat surveys conducted on 10 and 21 January 2026.

**Table S9: Summary of samples collected at Heard Island across October 2025 and January 2026 that were tested for Influenza A and high pathogenicity avian influenza (HPAI).** We endeavoured to sequence samples positive for HPAI. Where possible, swab samples were collected from the mouth, cloaca and brain for birds and mouth, nose and brain for seals. Fresh scat samples from live animals were collected and stored in ethanol.

| Species | Location | N animals swabbed | | | N scats | | | | N sequences (individuals) |
|---|---|---|---|---|---|---|---|---|---|
| | | Negative | HPAI | Total | Negative | Influenza A | HPAI | Total scats | |
| Southern elephant seal | Paddick Valley | | 21 | 21 | 2 | | | 2 | 18 |
| | Josephine Beach | | 3 | 3 | | | | | 3 |
| Antarctic fur seal | Skua beach | | 2 | 2 | | | | | 1 |
| Brown skua | Atlas Cove | | | | 1 | | | 1 | |
| | Josephine Beach | | | | 11 | | | 11 | |
| | Paddick Valley | | | | 2 | | | 2 | |
| | Skua Beach | | | | 8 | | 1 | 9 | |
| | North Spit | 1 | | 1 | | | | | |
| Gentoo penguin | Brown Lagoon | | | | 3 | | | 3 | |
| | Capsize Beach | | 1 | 1 | 10 | | | 10 | 1 |
| | Gauss Beach | | | | 4 | | | 4 | |
| | Skua Beach | | 1 | 1 | 16 | | | 16 | 1 |
| King penguin | Gauss Beach | | | | 1 | | | 1 | |
| | Josephine Beach | | | | 8 | | | 8 | |
| | Paddick Valley | | | | 7 | | 1 | 8 | 1 |
| | Schmidt Glacier | | | | 1 | | | 1 | |
| | Hoseason Beach | | | | 1 | | | 1 | |
| | North Spit | 4 | | 4 | | | | | |
| Macaroni penguin | Capsize Beach | | | | 10 | 1 | | 11 | |
| Sheathbill | Capsize Beach | | | | 1 | | | 1 | |
| Southern giant petrel | Brown Lagoon | | | | | 1 | | 1 | |
| | Gauss Beach | | | | 5 | | | 5 | |
| | North Spit | 1 | | 1 | | | | | |
| | Skua Beach | 1 | | 1 | | | | | |
| South Georgia diving petrel | Skua Beach | | 1 | 1 | | | | | |
| | **Total** | **7** | **29** | **36** | **91** | **2** | **2** | **95** | **25** |

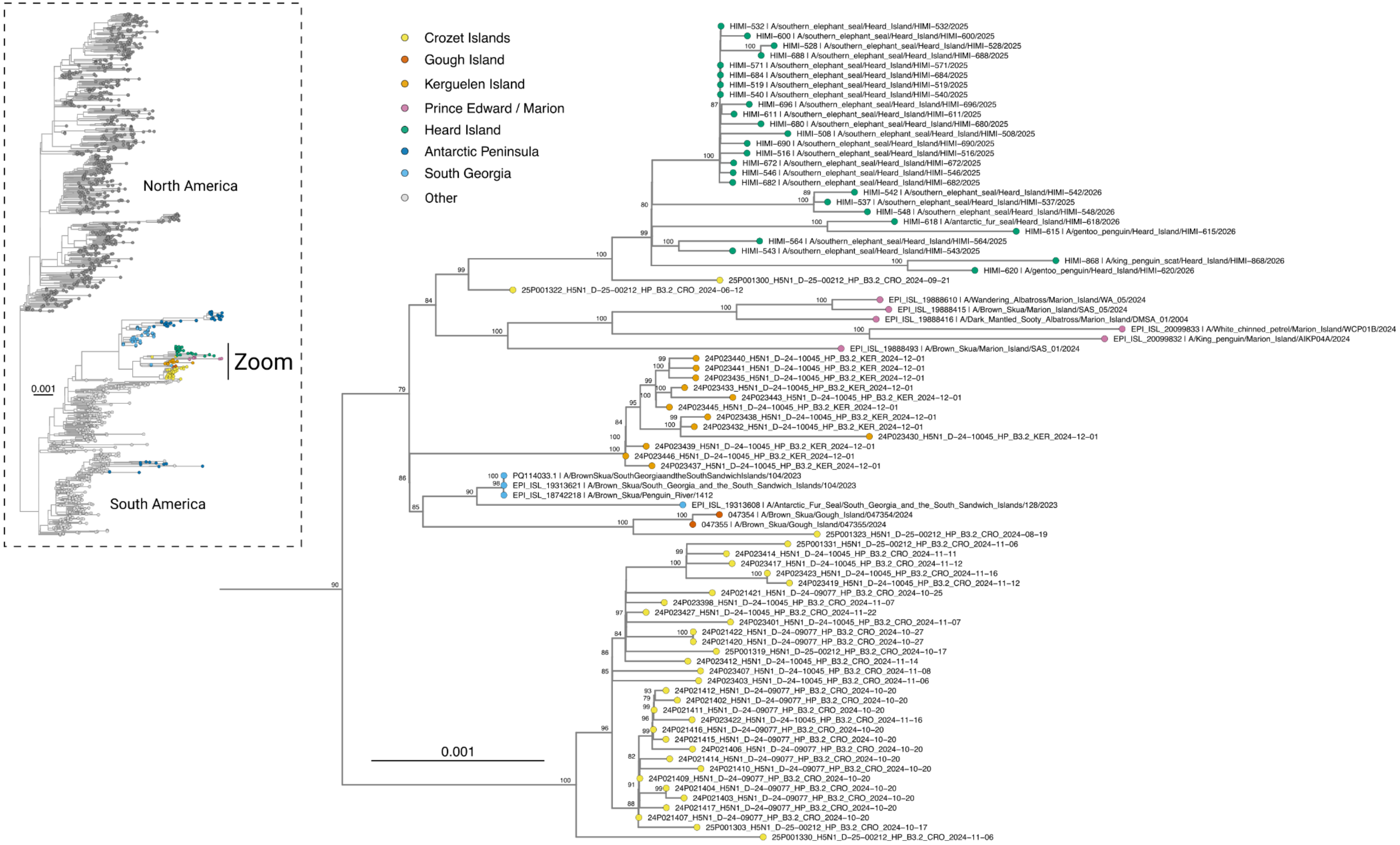

**Fig. S8: Clade I zoom of a maximum likelihood tree drawn from 1,322 concatenated genomes selected from North America, South America, Antarctica and sub-Antarctic islands (inset).** The scale indicates the average number of nucleotide substitutions per site and numbers on the nodes indicate percentages from 1,000 ultrafast bootstraps.

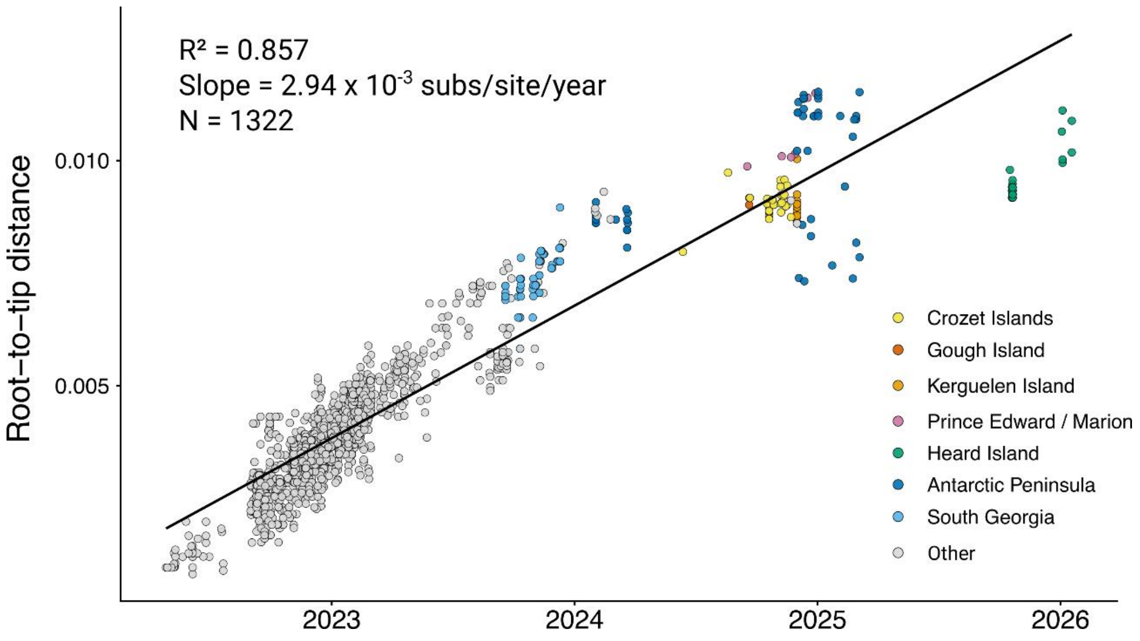

**Fig. S9: TempEst root-to-tip regression plot for 1,322 concatenated H5N1 2.3.4.4b whole genomes.** The regression slope, overall homoscedasticity and high $R^2$ value indicate strong temporal signal suitable for time-resolved phylogenetic analysis using an uncorrelated molecular clock model. Heard Island genomes had consistently negative residuals from the full-dataset regression line, with a mean residual of −0.00264 substitutions/site. Although the Heard Island cluster overall showed reduced root-to-tip divergence, the increase in genetic distance between viruses sampled in October 2025 and January 2026 appeared more consistent with the evolutionary rate estimated for the broader dataset ($2.94 \times 10^{-3}$ substitutions/site/year). However, this comparison is based on only two sampling periods and should be interpreted cautiously.

### REFERENCES

1 DotDotGoose (American Museum of Natural History, Center for Biodiversity and Conservation, Available from http://biodiversityinformatics.amnh.org/open_source/dotdotgoose, version 1.7.0).

2 Australian Antarctic Division, Department of External Affairs. Gibbney, L.F. *The Seasonal Reproductive Cycle of the Female Elepphant Seal – Mirounga leonina, Linn. – at Heard Island.* (1957).

3 Daniel, K., A. Nash, S. Koenig, & Felner, A. Theta*: Any-angle path planning on grids. *Journal of Artificial Intelligence Research* **39**, 533–579 (2010).

4 Wood, S. N. Generalized Additive Models. *Annual Review of Statistics and Its Application* **12**, 497-526 (2025).

5 Jocher, G. & Qiu, J. YOLO11 by Ultralytics https://github.com/ultralytics/ultralytics (2024).

6 Katoh, K., Rozewicki., J. & Yamada, K., D. MAFFT online service: Multiple sequence alignment, interactive sequence choice and visualization. *Briefings in Bioinformatics* **20**, 1160-1166 (2018).

7 Nguyen, L. T., Schmidt, H. A., Von Haeseler, A. & Minh, B.Q. IQ-TREE: A fast and effective stochastic algorithm for estimating maximum-likelihood phylogenies. *Molecular Biology and Evolution* **32**,268-274 (2015).

8 Rambaut, A., Lam, T. T., Carvalho, L. M & Pybus, O. G. Exploring the temporal structure of heterochronous sequences using TempEst (formerly Path-O-Gen). *Virus Evolution* **2** (2016).

9 McCann, T.S. Size, Status and Demography of Southern Elephant Seal (*Mirounga leonina*) Populations. in *Studies of Sea Mammals in South Latitudes* (eds. Ling, J.K. & Bryden, M.M.) 1-17 (South Australian Museum, 1985).
